# No changes in parieto-occipital alpha during neural phase locking to visual quasi-periodic theta-, alpha-, and beta-band stimulation

**DOI:** 10.1101/219766

**Authors:** Christian Keitel, Christopher SY Benwell, Gregor Thut, Joachim Gross

## Abstract

Recent studies have probed the role of the parieto-occipital alpha rhythm (8 – 12 Hz) in human visual perception through attempts to drive its neural generators. To that end, paradigms have used high-intensity strictly-periodic visual stimulation that created strong predictions about future stimulus occurrences and repeatedly demonstrated perceptual consequences in line with an entrainment of parieto-occipital alpha. Our study, in turn, examined the case of alpha entrainment by non-predictive low-intensity quasi-periodic visual stimulation within theta-(4 – 7 Hz), alpha-(8 – 13 Hz) and beta (14 – 20 Hz) frequency bands, i.e. a class of stimuli that resemble the temporal characteristics of naturally occurring visual input more closely. We have previously reported substantial neural phase-locking in EEG recording during all three stimulation conditions. Here, we studied to what extent this phase-locking reflected an entrainment of intrinsic alpha rhythms in the same dataset. Specifically, we tested whether quasi-periodic visual stimulation affected several properties of parieto-occipital alpha generators. Speaking against an entrainment of intrinsic alpha rhythms by non-predictive low-intensity quasi-periodic visual stimulation, we found none of these properties to show differences between stimulation frequency bands. In particular, alpha band generators did not show increased sensitivity to alpha band stimulation and Bayesian inference corroborated evidence against an influence of stimulation frequency. Our results set boundary conditions for when and how to expect effects of entrainment of alpha generators and suggest that the parieto-occipital alpha rhythm may be more inert to external influences than previously thought.

## INTRODUCTION

Human electroencephalographic (EEG) recordings contain prominent rhythmic components with an approximate 10 Hz periodicity, termed ‘alpha’ (Berger, 1929; Adrian & Matthews, 1934). Among them, the parieto-occipital alpha rhythm has received much attention because it has been consistently linked to perceptual processes (Bollimunta *et al.*, 2011; Clayton *et al.*, 2017). For example, parieto-occipital alpha power indexes the focus of visuo-spatial attention (Worden *et al.*, 2000; Kelly *et al.*, 2006; Thut *et al.*, 2006), allocation of intermodal attention (Banerjee *et al.*, 2011), or working memory load (Tuladhar *et al.*, 2007).

Pre-stimulus alpha power has also been linked to the perception of near-threshold transient stimuli (Busch & VanRullen, 2010; Benwell *et al.*, 2017b; Iemi *et al.*, 2017): Higher alpha power indicates lower cortical excitability and predicts decreased stimulus detection. Variations in alpha power have thus been proposed to reflect fluctuations in perceptual sensitivity (Linkenkaer-Hansen *et al.*, 2004; Busch *et al.*, 2009) or, more recently, shifts in response criterion and/or perceptual bias (Limbach & Corballis, 2016; Benwell *et al.*, 2017a; Benwell *et al.*, 2017b; Iemi *et al.*, 2017; Samaha *et al.*, 2017).

In parallel, a functional role has been ascribed to alpha phase: Cortical excitability waxes and wanes periodically within each alpha cycle (VanRullen *et al.*, 2014; VanRullen, 2016). Stimuli delivered during a specific phase of the cycle may thus experience cortical facilitation (or less inhibition), increasing their likelihood of being detected, relative to stimuli presented during the opposite phase (Mathewson *et al.*, 2009). This process can be adaptive: Transient alpha frequency changes produce phase shifts that serve to accommodate expected upcoming stimulation during optimal alpha phase, i.e. periods of relatively high cortical excitability (Samaha & Postle, 2015).

Based on the notion that alpha phase indexes fluctuations in cortical excitability and, consequentially, perceptual outcome, attempts have been made to interact with pre-stimulus alpha activity to influence visual target processing. Recent findings from electrophysiology suggest that alpha phase can synchronize to continuous, rhythmic visual stimulation (Spaak *et al.*, 2014; Notbohm *et al.*, 2016; Gulbinaite *et al.*, 2017);(but see Capilla *et al.*, 2011; Keitel *et al.*, 2014). In terms of behavioural consequences, this has been associated with a greater chance of transient targets being detected when they occur in-phase with the ongoing (Sokoliuk & VanRullen, 2016) or just-ceased alpha-rhythmic visual stimulation (Mathewson *et al.*, 2012; de Graaf *et al.*, 2013; Spaak *et al.*, 2014). Put differently, targets were detected more often when they occurred at a time at which a stimulus presentation could be expected.

The above findings on alpha phase synchronisation agree with model predictions of how a population of self-sustained neuronal oscillators could *entrain* to a rhythmic external drive (Thut *et al.*, 2011; also see Herrmann *et al.*, 2016a). For these effects to be explained in an entrainment framework, the following requirements (amongst others) need to be met: (1) Neuronal oscillators must be in place to be able to follow the (periodic) temporal structure of the visual input, eventually leading to a phase synchronisation. Regarding alpha-rhythmic visual stimulation (in the ∼10 Hz range), this requirement is easily met because alpha rhythms dominate visual cortex activity (Keitel & Gross, 2016) and the visual cortex response to transcranial magnetic stimulation resonates particularly strongly at its natural eigenfrequencies (Rosanova *et al.*, 2009; Herring *et al.*, 2015). (2) The external drive needs to be periodic – a feature that has recently been supported experimentally by comparing strictly periodic to irregular stimulation (Notbohm & Herrmann, 2016).

The model further identifies influential factors, one of which is the strength of the external drive (Pikovsky *et al.*, 2003). For visual stimulation a strong drive can be quantified as a high peak-to-peak change in luminance or contrast. Another factor, recently suggested to play a central role especially in visual perception, is the task-relevance of the entraining stimulation (Zoefel & VanRullen, 2017).

To date, most studies into the entrainment of endogenous alpha have indeed employed high-intensity, strictly periodic stimulation that was task-relevant, i.e. the stimulus phase generated predictions about the occurrence of the target (Mathewson *et al.*, 2012; de Graaf *et al.*, 2013; Spaak *et al.*, 2014; Notbohm & Herrmann, 2016). However, the limiting factors under which entrainment of endogenous alpha occurs or ceases have yet to be examined. This will be necessary in order to comprehensively characterise it as a fundamental neural process that may contribute to visual perception. To date, it remains an open question whether alpha entrainment occurs automatically in natural vision because continuous, ecologically valid stimulation is quasi-periodic at best, does not need to be of high contrast and does not necessarily fall within the alpha frequency range (Kayser *et al.*, 2003; Blake & Lee, 2005), e.g. lip movements during speech (Chandrasekaran *et al.*, 2009; Park *et al.*, 2016).

In the present report, we thus tested whether quasi-periodic contrast modulations of low-intensity visual input produce effects consistent with an entrainment of endogenous alpha rhythms. Data from a recent EEG study allowed us to evaluate the influence of quasi-periodic stimulation within three distinct frequency bands – theta (4-7 Hz), alpha (8-13 Hz) and beta (14-20 Hz) – on properties of the intrinsic alpha rhythm and its functional characteristics (Keitel *et al.*, 2017c). Note that our stimulation was not immediately task-relevant in the sense that stimulus phase had no predictive value for target detection: Participants viewed two stimuli (see *Figure 1*), one in each visual hemifield, that underwent concurrent quasi-periodic contrast modulations. They were cued to attend to the location of one of the two stimuli, positioned in lower left and right hemifields, and performed a visual detection task.

**Figure 1.**
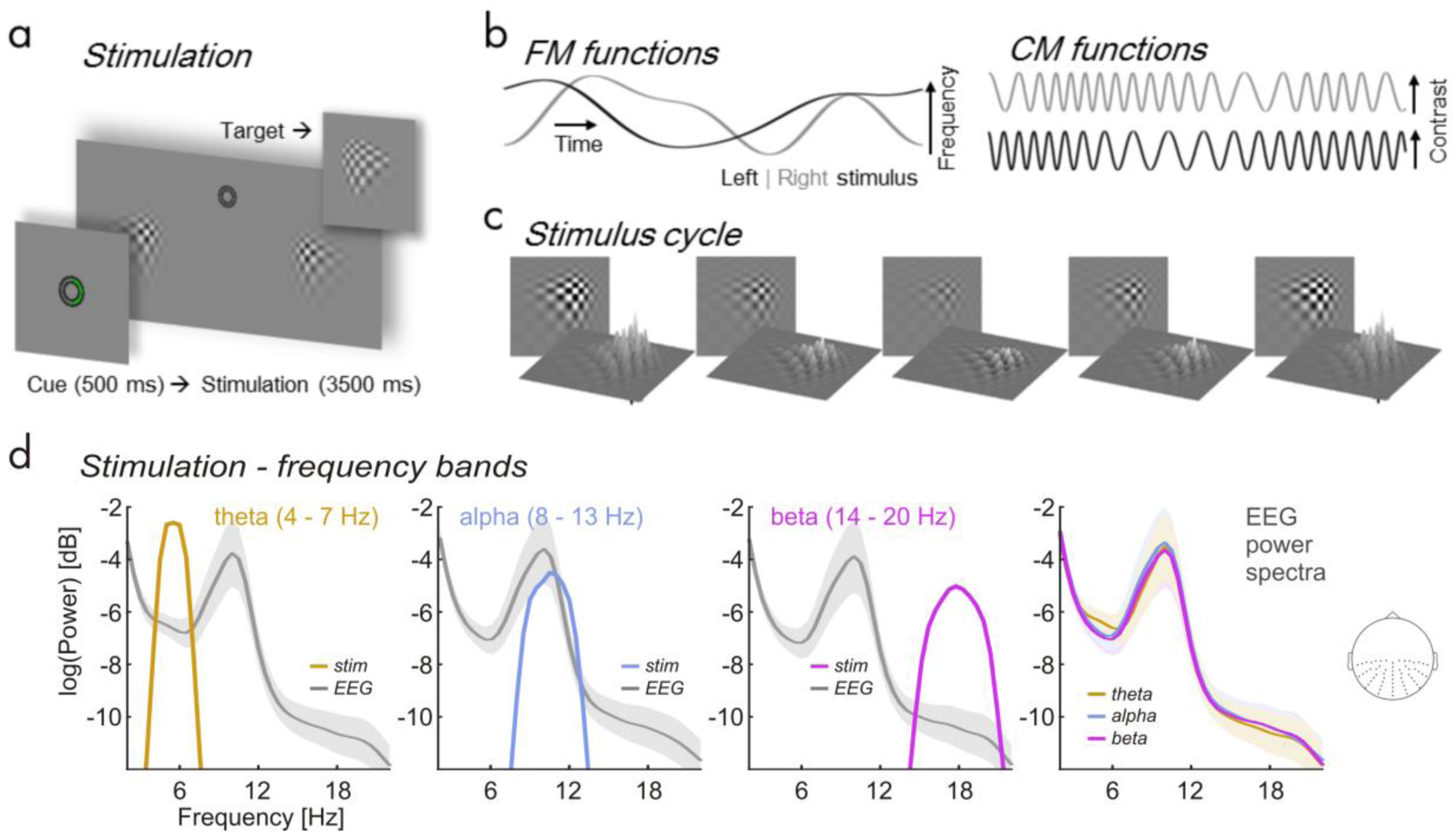
Details of the experimental stimulation. (**a**) Trial time course. Central cue presentation (green semi-circle, “attend right”) precedes continuous streams of contrast modulating patches. Upper right inset gives an example of target (and distracter) appearances. (**b**) Time series depict random band-limited frequency fluctuations (FM functions, left graphs) in periodic contrast modulation functions (CMFs, right graphs) for left and right stimuli on a given trial. (**c**) Left stimulus traversing one CMF peak-to-peak cycle. (**d**) Spectral profiles of band-limited stimuli within theta (orange), alpha (blue) and beta frequency ranges (purple) and corresponding grand average (N = 17) EEG power spectra during stimulation (grey). Note that a constant was added to stimulus power spectra to match the scale of EEG power spectra for illustrative purposes. Right-most plot: Overlay of EEG power (logarithmic scale) during theta-, alpha- and beta band visual stimulation. Spectra were pooled across electrodes indicated on the inset scalp map. Shaded areas around EEG power spectra depict standard error of the mean.

Our previous EEG analysis revealed that brain responses elicited by these low-intensity stimuli followed their temporal evolution (*Figure 1b*), thereby suggesting a neural phase synchronisation^1^ (Keitel *et al.*, 2017c). Quantifying this synchronisation required calculating a measure of phase locking between the frequency-varying stimulation and the EEG (Gross *et al.*, 2013; Peelle *et al.*, 2013). A spectral representation of EEG-stimulus locking showed clear peaks within the stimulated frequency ranges. While analysing these data, we noticed that EEG power spectra were virtually unaffected by the stimulation but instead were dominated by the typical alpha peak (see *Figure 1d*). This suggested the absence of an influence of rhythmic visual input on the generation of intrinsic alpha rhythms in any stimulation condition (theta, alpha, & beta). This observation prompted the present, detailed investigation of the functional and systemic characteristics of the intrinsic alpha rhythm under the different stimulation conditions. More specifically, we re-examined these data for indicators of visual entrainment of endogenous alpha. We focussed on the typical retinotopic alpha lateralisation effect by attention (Kelly *et al.*, 2006; Thut *et al.*, 2006), as enabled by our spatial attention manipulation, and tested whether it depended on the frequency of the stimulation. We further tested whether intrinsic alpha resonates during alpha-band stimulation (Fedotchev *et al.*, 1990; Schwab *et al.*, 2006; Spiegler *et al.*, 2011), i.e. shows relatively greater power increase than theta power during theta-band stimulation and beta power during beta-band stimulation. Lastly we probed individual alpha peak frequency for shifts towards the stimulation’s centre frequency as would be expected during alpha entrainment.

## METHODS

### Participants

For the present report, we re-analysed EEG data and behavioural performance of 17 volunteers recorded in an earlier study (Keitel *et al.*, 2017c)^2^. Participants (13 women; median age = 22 yrs, range = 19 – 32 yrs) declared normal or corrected-to-normal vision and no history of neurological diseases or injury. All procedures were approved by the ethics committee of the College of Science & Engineering at the University of Glasgow (application no. 300140020). Volunteers received monetary compensation. They gave informed written consent before participating in the experiment.

### Stimulation

Participants viewed experimental stimuli on a computer screen (refresh rate = 100 frames per sec) at a distance of 0.8 m and displaying a grey background (luminance = 6.5 cd/m^2^). Small concentric circles in the centre of the screen served as a fixation point (*Figure 1a*). At an eccentricity of 4.4° from central fixation, two blurry checkerboard patches (horizontal/vertical diameter = 6°/4.4° of visual angle) were positioned, one each in the lower left and lower right visual quadrants. Both patches changed contrast dynamically during trials: Stimulus contrast against the background was modulated by varying patch peak luminance between 7.5 cd/m^2^ (minimum) and 29.1 cd/m^2^ (maximum). On each screen refresh peak luminance changed incrementally to approach temporally smooth contrast modulations (*Figure 1b and c*) as opposed to a simple on-off flicker (Andersen & Muller, 2015). Further details of the stimulation can be found in Keitel *et al.* (2017c).

Contrast modulations were generated for each experimental trial and both patch stimuli independently and obeyed one of three different spectral profiles (*Figure 1d*). In trials of the theta-rhythmic condition, contrast modulation rates varied between 4 – 7 Hz, in the alpha-rhythmic condition between 8 – 13 Hz and in the beta-rhythmic condition between 14 – 20 Hz. Frequencies were limited to maximally change by one bandwidth per second (e.g. ∼3 Hz/sec for theta-rhythmic stimulation). Frequency modulations were controlled to be maximally uncorrelated between concurrently presented stimuli (Pearson correlation coefficient *r* < 0.05). Note that the experiment further featured a condition in which stimulus contrast was modulated at constant rates of 10 and 12 Hz. Corresponding data are reported in the original study and will not be considered in the primary analyses – for an exemplary investigation of alpha functional modulation during strictly rhythmic stimulation see Kelly *et al.* (2006). This exclusion is based on the fact that the constant-frequency stimulation differs in more than one aspect (strictly rhythmic, consistently distinct frequencies left and right) from the frequency-varying conditions. Therefore, statistical differences arising between conditions cannot be unambiguously attributed to either manipulation.

This caveat notwithstanding, the Supplementary Material contains a section *Analyses including the strictly-rhythmic stimulation condition* that retraces all analysis steps described below including the constant-frequency stimulation conditions for future reference.

### Procedure and Task

Participants performed the experiment in an acoustically dampened and electromagnetically shielded chamber. In total, they were presented with 576 experimental trials, subdivided into 8 blocks with durations of ∼5 min each. Between blocks participants took self-paced breaks. Prior to the experiment, participants practiced the behavioural task (see below) for at least one block. After each block they received feedback regarding their accuracy and response speed. The experiment was comprised of 8 conditions (= 72 trials each) resulting from a manipulation of the two factors *attended position* (left vs. right patch) and *stimulation frequency* (constant, theta, alpha and beta) in a fully balanced design. Trials of different conditions were presented in pseudo-random order. As stated above, here we focus on the frequency-varying conditions of the experiment. Corresponding trials were thus selected a posteriori from the full design. (For an analysis including the constant-frequency stimulation conditions see section *Analyses including the strictly-rhythmic stimulation condition* of the Supplementary Material.)

Single trials began with cueing participants to attend to the left or right stimulus (*Figure 1a*) for 0.5 sec, followed by presentation of the dynamically contrast-modulating patches for 3.5 sec. After patch offset, an idle period of 0.7 sec allowed participants to blink before the next trial started.

To control whether participants maintained a focus of spatial attention they were instructed to respond to occasional brief “flashes” (0.3 sec) of the cued stimulus (= targets) while ignoring similar events in the other stimulus (= distracters). Targets and distracters occurred in one third of all trials and up to 2 times in one trial with a minimum interval of 0.8 sec between subsequent onsets. Detection was reported as speeded responses to flashes (recorded as space bar presses on a standard key board).

### Behavioural data recording and analyses

Flash detections were considered a ‘hit’ when a response occurred from 0.2 to 1 sec after target onset. Delays between target onsets and responses were considered reaction times (RT). Statistical comparisons of mean accuracies (proportion of correct responses to the total number of targets and distracters) and median RTs between experimental conditions were conducted and reported in Keitel *et al.* (2017c). In the present study, we further tested how individual alpha power influenced task performance based on these data as described below.

### Electrophysiological data recording

EEG was recorded from 128 scalp electrodes and digitally sampled at a rate of 512 Hz using a BioSemi ActiveTwo system (BioSemi, Amsterdam, Netherlands). Scalp electrodes were mounted in an elastic cap and positioned according to an extended 10-20-system (Oostenveld & Praamstra, 2001). Lateral eye movements were monitored with a bipolar outer canthus montage (horizontal electro-oculogram). Vertical eye movements and blinks were monitored with a bipolar montage of electrodes positioned below and above the right eye (vertical electro-oculogram).

### Pre-processing

#### Electrophysiological data during stimulation

From continuous data, we extracted epochs of 5 s starting 1 s before patch onset (= cue offset) using the MATLAB toolbox EEGLAB (Delorme & Makeig, 2004). In further pre-processing, we excluded epochs that corresponded to trials containing transient targets and distracters (24 per condition) as well as epochs with horizontal and vertical eye movements exceeding 20 μV (∼ 2° of visual angle) or containing blinks. For treating additional artefacts, such as single noisy electrodes, we applied the ‘fully automated statistical thresholding for EEG artefact rejection’ (FASTER, Nolan *et al.*, 2010). This procedure corrected or discarded epochs with residual artefacts based on statistical parameters of the data. Artefact correction employed a spherical-spline-based channel interpolation. Epochs with more than 12 artefact-contaminated electrodes were further excluded from analysis.

From 48 available epochs per condition, we discarded a median of 12 epochs (25 %) per participant with a between-subject range of 5 to 20.5 epochs (10 – 43%). Because within-participant variation of numbers of epochs per condition remained small (with a median maximum difference of 8 trials between conditions) trial numbers were not artificially matched.

Subsequent analyses were carried out in Fieldtrip (Oostenveld *et al.*, 2011) in combination with custom-written routines. We extracted segments of 3 s starting 0.5 s after patch onset from pre-processed artefact-free epochs (5 s). Data prior to stimulation onset (1 s), only serving to identify eye movements shortly before and during cue presentation, were omitted. To attenuate the influence of stimulus-onset evoked activity on EEG spectral decomposition, the initial 0.5 s of stimulation were excluded. Lastly, because stimulation ceased after 3.5 s, we also discarded the final 0.5 s of original epochs. The remaining data were further segmented into epochs of 1 s with an overlap of 0.5 s.

#### Electrophysiological data peri-stimulation

In addition to data allowing EEG analyses during stimulation, we extracted epochs of 6 sec immediately *before* experimental blocks started and *after* blocks ended to obtain a measure of the participants’ alpha rhythm shortly before and after (‘peri-’) stimulation using Fieldtrip (Oostenveld *et al.*, 2011). Participants were instructed to remain still during these periods. Before blocks they viewed a 6-sec countdown preparing them for the upcoming stimulation and after blocks they were prompted to patiently expect the processing of their behavioural track record for a similar duration. Data from the resulting 16 segments was concatenated and cut into epochs of 1 sec with an overlap of 0.5 sec. Artefact-contaminated epochs and channels were rejected by visual inspection (fieldtrip function *ft_rejectvisual*, method ‘*summary’*). For each participant, we retained more than 100 epochs (range: 102 – 171, median = 134) and excluded up to 4 channels (median = 2). After re-referencing the data to average reference, an independent component analysis using the logistic infomax algorithm (Bell & Sejnowski, 1995) served to minimize contamination by blinks and eye movements: Per participant, between 1 and 3 independent components were removed. Globally contaminated channels, removed during visual inspection, were spherical-spline interpolated after component removal.

Note that an ICA for eye artefact removals was not applied on EEG data recorded during stimulation. Instead trials containing eye artefacts were simply discarded because blinks interrupted the continuous visual input and eye movements may have changed retinotopic projections to visual cortex, greatly influencing the stimulus-driven brain response (Walter *et al.*, 2012).

### Electrophysiological data analyses

EEG data analyses were aimed at identifying modulations of the intrinsic alpha rhythm as a function of stimulation frequency. Different analysis paths are illustrated in Figure 2 and laid out in detail below. In brief, we focussed on the following aspects of the data: First we sought to replicate the typical pattern of hemispheric alpha power lateralisation observed during endogenous shifts of spatial attention to the left or right hemifield positions (Kelly *et al.*, 2006; Thut *et al.*, 2006) and compared attentional modulations between conditions. This step further served to define clusters of EEG sensors on which to conduct subsequent analyses.

**Figure 2.**
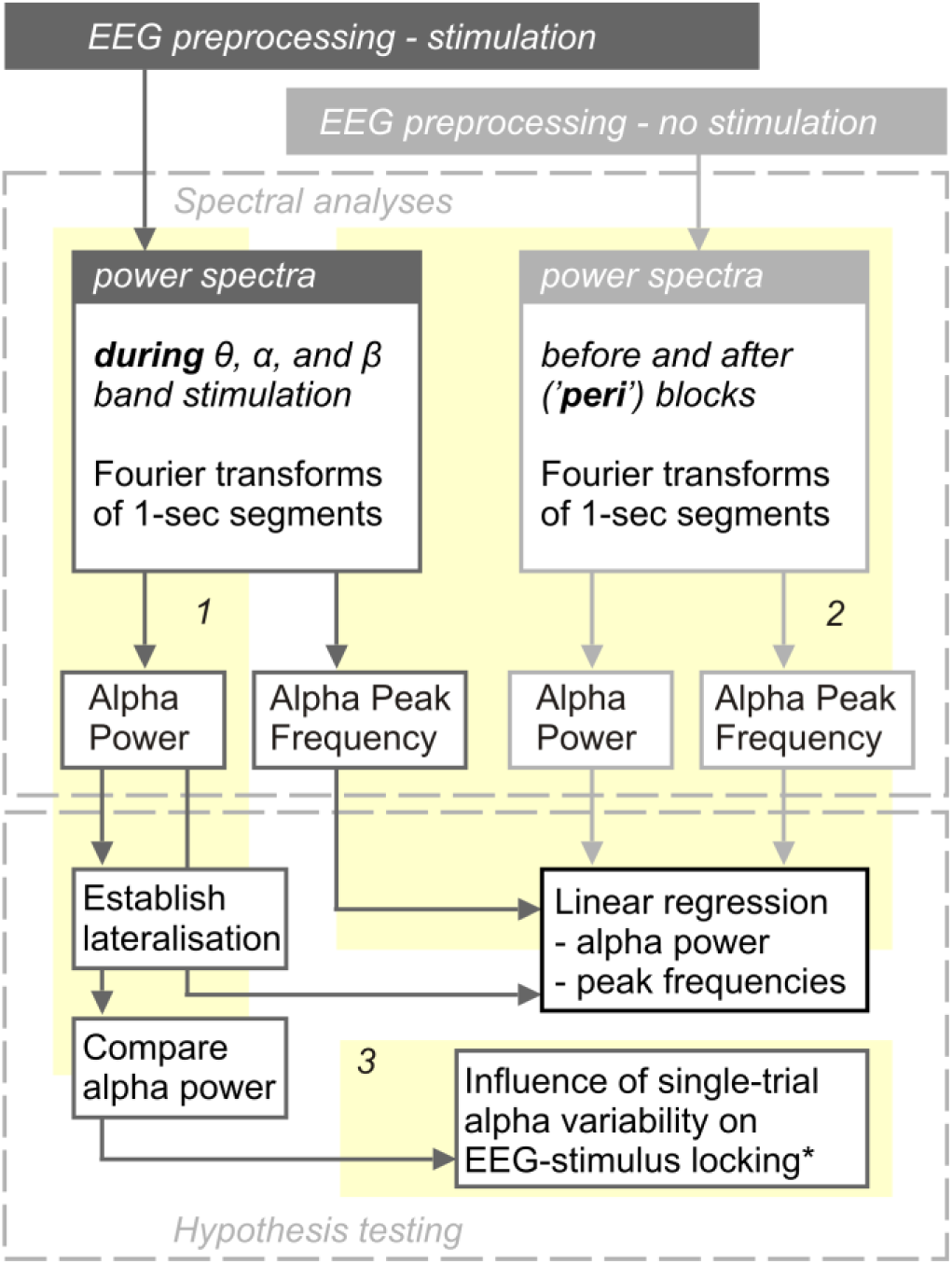
Analysis flow. Each participants EEG recording was preprocessed differently depending on whether they received visual stimulation or not (i.e. shortly before and after experimental blocks = ‘peri’-stimulation). Preprocessed data entered three main analyses paths (numbered yellow backdrops *1, 2* and *3*). Path 1 and 2 extracted individual alpha power and peak frequencies. Path 1 investigated retinotopic alpha power lateralisation during focused spatial attention, allowed a comparison of alpha power between stimulation conditions and provided input for Path 3. Alpha power and peak frequencies extracted along Path 2 were used to regress similar parameters extracted along Path 1. Detailed methods and results regarding Path 3 are documented in the *Supplementary Material*. * EEG-stimulus locking analyses are described in detail in Keitel et al. (2017c).

Secondly, we correlated alpha power and individual alpha peak frequency (IAF) during stimulation with the same measures peri-stimulation. Data obtained in this step also served to establish links between individual alpha power during stimulation and behavioural performance.

In an additional analysis, we explored single-trial variations in alpha power lateralisation at individual peak frequencies and their influence on neural phase-locking to the visual stimulation (see *Supplementary Material, Figures S2 & S3*).

All EEG analyses were conducted on frequency-domain data obtained as described in the following.

#### Spectral decomposition

Epochs of 1 s length, derived from EEG data during-as well as peri-stimulation, were converted to scalp current densities (SCDs), a reference-free measure of brain electrical activity (Ferree, 2006; Kayser & Tenke, 2015), by means of the spherical spline method (Perrin *et al.*, 1987) as implemented in Fieldtrip (function *ft_scalpcurrentdensity*, method ‘spline’, lambda = 10^-4^). Detrended (i.e. mean and linear trend removed) SCD time series were then Hanning-tapered and subjected to Fourier transforms while employing zero-padding in order to achieve a frequency-resolution of 0.5 Hz. We calculated power spectra as the squared absolute values of complex Fourier spectra. IAF was determined based on raw power spectra (see section *Alpha power and peak frequency during vs peri-stimulation* below). For further analyses, power spectra were normalised by converting them to decibel scale, i.e. taking the decadic logarithm, then multiplying by 10 (hereafter termed *log-power* spectra).

#### Alpha power attentional modulation and lateralisation

We identified two sets of parieto-occipital electrodes, one above each hemisphere, which showed systematic variation in alpha power by attentional allocation as follows: Mean log-power across the range of 8 – 12 Hz was pooled over conditions in which participants either attended to the right or left stimulus positions separately yielding two alpha power topographies for each participant. These were compared by means of cluster-based permutation statistics (Maris & Oostenveld, 2007) using *N* = 1000 random permutations. We clustered data across channel neighbourhoods with an average size of 7.9 channels that were determined by proximity in a planar projection of 3D electrode locations (function *ft_prepare_neighbours*, method ‘triangulation’). The resulting probabilities (*P*-values) were corrected for two-sided testing.

We expected to find a right-hemispheric positive and a left-hemispheric negative cluster of electrodes because we subtracted right-lateralized (Attend-Right conditions) from left-lateralized (Attend-Left) alpha power topographies. This approach yielded two functionally defined left and right parieto-occipital clusters (see *Figure 3*) that were used in all subsequent analyses.

**Figure 3.**
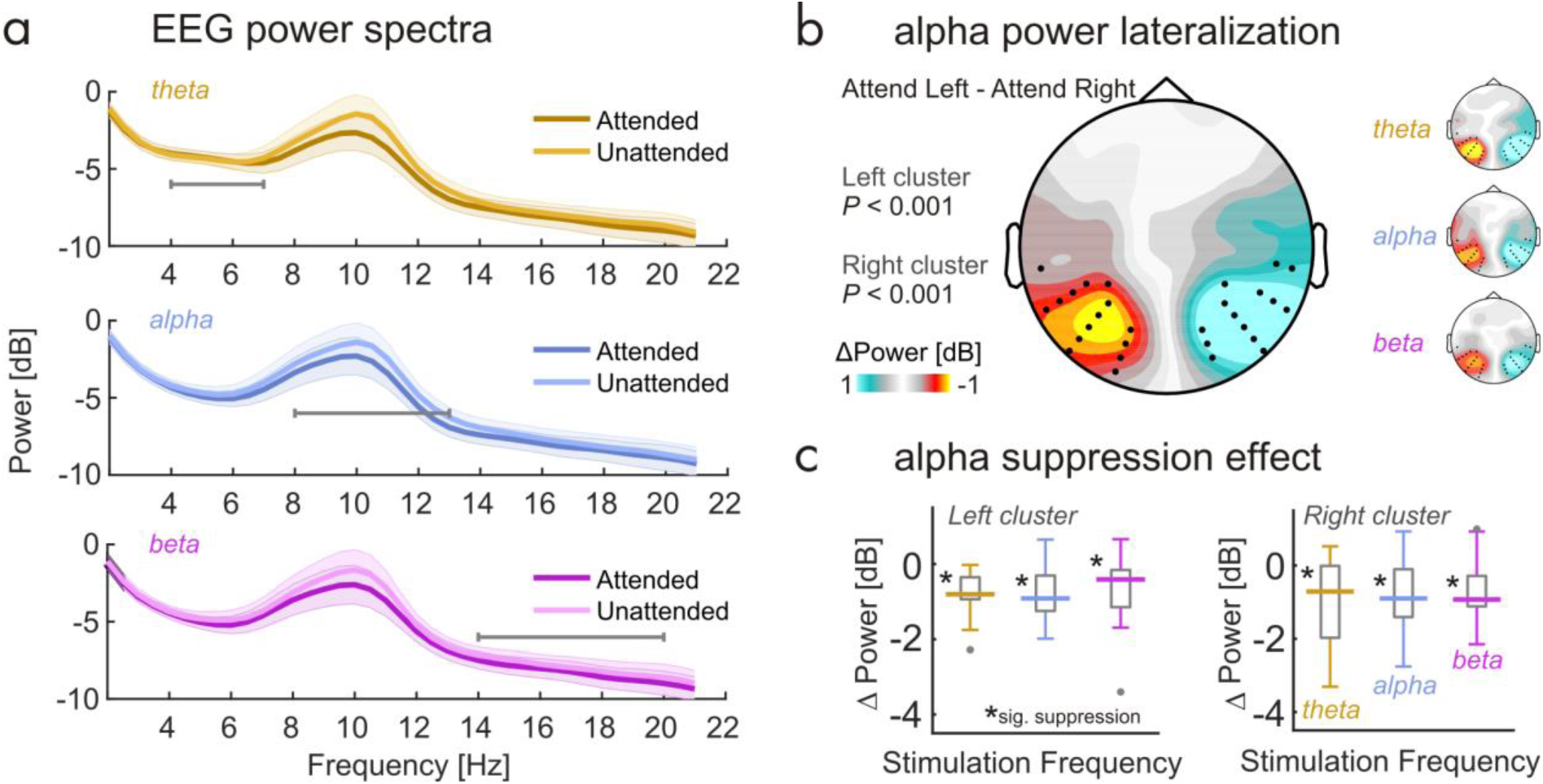
Attentional allocation produces alpha lateralisation. (**a**) Log-power spectra during theta-band (golden), alpha-band (blue) and beta-band (purple) stimulation collapsed across electrode clusters depicted in (b) and aggregated across ‘Attend Left’ and ‘Attend Right’ conditions to reveal the attention effect on alpha power. Note the prominent difference in the range 8 - 12 Hz. Shaded errors show pointwise standard error of the mean. Horizontal grey lines indicate respective stimulated frequency ranges. (**b**) Scalp map depicts alpha lateralisation collapsed across stimulation conditions during spatially focussed attention. Warm colors signify lateralisation of alpha power when subjects focus towards the right hemifield. Cool colors signify focus towards the left hemifield position. Black dots indicate two clusters of electrodes that demonstrated significant lateralisation (both *P* < .001, cluster corrected) and that were used in further analyses. Small inset scalp maps on the right illustrate condition specific alpha lateralisation. (**c**) Boxplots depicting distributions of alpha suppression effects (Attended minus Unattended alpha power) for the three stimulation frequency bands and for left and right hemispheric electrode clusters (see b), separately. Boxes cover the interquartile range, strong horizontal lines signify medians. Grey dots represent outlier. Asterisks near median lines demarcate significant suppression on a Holm-Bonferroni corrected alpha level (*P*_HB_ < .05).

As for average alpha power within the 8-12 Hz band, we conducted further statistical analyses. A three-way repeated-measures analysis of variances (ANOVA) probed differences in alpha power between *hemispheres* (left vs right electrode clusters), attended *stimulus positions* (contra-vs ipsilateral hemifield, with respect to *hemisphere*) and *stimulation frequency* (theta, alpha or beta). In all repeated measures ANOVAs conducted in this study that featured factors with more than two levels, the Greenhouse–Geisser (GG) adjustment of degrees of freedom was applied to control for violations of sphericity (Greenhouse & Geisser, 1959). Original degrees of freedom, corrected p-values (*P*_GG_) and the correction coefficient epsilon (ε_GG_) are reported for significant effects. Effect sizes are documented as *ω*_*p*_^2^ (Keren & Lewis, 1979). Further specific contrasts of alpha power modulations (attended minus unattended) against zero employed two-tailed t-tests. The resulting *P*-Values were Holm-Bonferroni-corrected (reported as *P*_HB_) for multiple comparisons (Holm, 1979).

In addition to the classical frequentist approach described above, we also applied Bayesian inference. To this end, we estimated Bayes factors (Bf) in R (R-Core-Team, 2016) by means of the function *anovaBF* provided with the package *Bayes factor* (Morey & Rouder, 2015) running in RStudio (RStudio-Team, 2015). We adopted the Jeffrey-Zellner-Siow (JZS) prior with a scaling factor *r* of .5 that puts more emphasis on smaller effects (Rouder *et al.*, 2009; Rouder *et al.*, 2012; Schonbrodt & Wagenmakers, 2017). Bf estimation involved Monte-Carlo resampling with 10^5^ iterations. Participants were considered as a random factor.

Crucially, this Bayesian equivalent to an ANOVA allowed us to quantify the evidence that our experimental factors (*hemisphere, attended position* and *stimulus frequency*), as well as their interactions, explained variance in the data beyond the random between-subject fluctuations. In other words, we tested whether we have more evidence in favour of a factor having no systematic influence (null hypothesis, *H*_*0*_) on alpha power vs having an influence (alternative hypothesis, *H*_*1*_). For each test, the corresponding Bayes factor (called BF_10_) shows evidence for H_1_ (compared to H_0_) if it exceeds a value of 3, and no evidence for H_1_ if BF_10_ < 1, with the intervening range 1 – 3 termed ‘anecdotal evidence’ by convention (Wagenmakers *et al.*, 2011). Inversing BF_10_, to yield a quantity termed BF_01_, serves to quantify evidence in favour of H_0_ on the same scale. For BF_10_ and BF_01_ values < 1 are taken as inconclusive evidence for either hypothesis.

#### Alpha power and peak frequency during vs peri-stimulation

To test for stimulation-specific effects on alpha-rhythmic activity, we compared two characteristics, individual alpha peak frequency (IAF) and alpha power between EEG data recorded during stimulation and data recorded shortly before and after stimulation.

First, we determined IAF based on raw power spectra for peri-stimulation data, as well as for the three stimulation conditions separately. To this end, we took the second order gradient of power spectra averaged across the two electrode clusters (as extracted in stage 1). Sign-flipped (i.e. multiplied by −1) gradient spectra were then smoothed by means of an 11-point, third-order polynomial Savitzky-Golay filter (Savitzky & Golay, 1964). This procedure was implemented to accentuate peaks in participants with indistinct maxima in raw spectra while providing virtually identical results for participants with clear alpha peaks (see *Figure S1*). A similar step has been implemented in a recent algorithmic approach of IAF determination (Corcoran *et al.*, 2017).

Individual alpha power (IAP) was extracted as the average spectral power within the band IAF ± 1 Hz from log-power spectra averaged across electrodes. Using IAFs, we further derived a measure of individual alpha lateralisation, for which we subtracted IAP collapsed across sensors of the cluster ipsilateral to the attended position from IAP collapsed across contralateral sensors in each condition.

First, we tested whether peri-stimulation IAF/IAP predicted measurements during stimulation by means of regressions using robust linear fits (Matlab function *robustfit*) for all three stimulation conditions separately. Linear fits are determined by two coefficients, their intercept (*β*_*0*_) and their slope (*β*_*1*_). The intercept did not have a meaningful interpretation for IAF. We thus standardized (z-scored) IAF across participants prior to regression to focus on the slope as a possible indicator of entrainment of endogenous alpha. In the case of individual alpha power, the intercept (*β*_*0*_) could have been an indicator of a general power increase (or decrease) during stimulation. We thus ran regression analyses on non-standardized log-power values.

Statistical significance of regression coefficients was assessed by bootstrapping (*N* = 1000) two-tailed 95% confidence intervals (CIs) involving the bias-corrected and accelerated percentile method (Efron, 1987) to control for deviations from normality of the underlying bootstrapped distribution. The null hypothesis (H_0_) was rejected when the bootstrapped CI excluded zero.

Subsequently, when testing for systematic differences *between* slopes (i.e. between conditions), H_0_ was rejected when the CI based on the bootstrapped distribution of slope *differences* excluded zero. Due to the post-hoc nature of these tests, CIs were evaluated on a Bonferroni-adjusted significance level of 1-0.05/n (hereafter termed *CI*_*BF*_) with n = 3, i.e. the maximum number of pairwise comparisons within each family of regressions.

Employing a similar approach, we explored the relationship between individual alpha power during stimulation and behavioural performance in the visual detection task. To this end, we transformed the performance measure (proportion of correct responses (in %)) using a logit-link function as a means of normalisation and removing bounds. Then we tested whether standardized (z-scored) IAP, as well as alpha lateralisation during theta, alpha and beta-band stimulation predicted respective standardised performance measures using robust linear regression. In case of yielding condition-specific significant regression slopes, these were subjected to pairwise comparisons between conditions as described above.

## RESULTS

### Attention modulates alpha power retinotopically

Attention modulated alpha power as expected (Figure 3a). In line with previous findings, cluster-based permutation statistics revealed two parieto-occipital electrode clusters (both *P* < .001) at which alpha power (8 – 12 Hz) systematically varied between Attend-right and Attend-left conditions (*Figure 3b*). In both cases, alpha power decreased when participants attended to a stimulus position in the contralateral hemifield, thereby mirroring the known signature of lateralised attention orienting. Scalp maps of condition-specific alpha power contrasts (Attend Left minus Attend Right) in *Figure 3b* closely resembled the condition-collapsed contrasts.

Grand average log-power spectra in *Figure 3a* illustrate alpha suppression effects when participants attended contralateral hemifields (collapsed across left and right electrode clusters) in all three stimulation conditions. Testing these differences (Attended minus Unattended) against zero separately for left and right clusters yielded substantial modulations of alpha power for each stimulation frequency (all six comparisons: *P*_HB_ < .05, see *Figure 3c*).

### No evidence for an influence of visual stimulation frequency on alpha power

A repeated-measures ANOVA based on the data pooled within each cluster confirmed a general decrease in alpha power when the contralateral stimulus position was attended (main effect *attended position*: *F*(1,16) = 43.24, *P* < 0.001, *ω*_*p*_^2^ = 0.701; also see *Figure 3c*). Alpha power was comparable in the left and right clusters (main effect *hemisphere*: *F*(1,16) = 3.20, *P* = 0.092, *ω*_*p*_^2^ = 0.109). Crucially however, the frequency composition of the visual stimulation did not influence alpha power (main effect *stimulus frequency*: *F*(2,32) = 1.96, *P* = 0.158, *ω*_*p*_^2^ = 0.052) and factor interactions were not significant (*F*_max_(1,16)= 1.95, *P*_min_ = 0.162).

Bayesian inference also showed strong evidence for an influence of *attended position* on alpha power (BF_10_ = 4,195,405.000 ± 0.55%). Also, we found strong evidence for an imbalance in overall alpha power between hemispheres (BF_10_ = 171.166 ± 0.69%) that the frequentist ANOVA did not capture. In fact, alpha power measured over the right hemisphere (*M* = −2.807 dB, *SEM* = ± 1.022) was greater than alpha power measured over the left hemisphere (*M* = −3.340 dB, *SEM* = ± 1.029). However, and in line with the above results, the factor *stimulus frequency* showed substantial evidence in favour of *H*_*0*_, i.e. no effect of stimulation frequency on alpha power (1/BF_10_ = BF_01_ = 5.962 ± 0.26%). The remaining factors and interactions failed to provide conclusive evidence for *H*_*1*_ or *H*_*0*_ (all BF_10_ < 0.22).

Note that the absence of an effect of alpha-band stimulation on alpha power precluded further dedicated analyses of an alpha resonance effect, i.e. a greater increase in alpha power during alpha band stimulation than in theta power during theta-band or beta-power during beta-band stimulation. Although our stimulation may have produced increased theta power (see Figure 1d, right-most panel), we did not find a similar effect in the alpha band to compare this increase to.

### No evidence for interaction of frequency of visual stimulation with power and peak frequency of intrinsic alpha oscillations

We hypothesised that intrinsic alpha rhythms should be most effectively entrained during alpha-band stimulation. One first indicator of pronounced entrainment could have been a resonance phenomenon, expressed as a greater increase in alpha power during alpha-band stimulation than during theta- or beta-band stimulation compared with alpha power when no stimulation was presented (Fedotchev *et al.*, 1990; Schwab *et al.*, 2006; Herrmann *et al.*, 2016a).

Secondly, assuming that alpha generators were indeed contributing to the neural response to the ongoing stimulation, we expected individual alpha frequencies, measured in the absence of stimulation, to be pulled towards the centre frequency of the alpha band stimuli during stimulation (Thut *et al.*, 2011; Herrmann *et al.*, 2016b). This effect of entraining endogenous alpha has recently been demonstrated with transcranial alternating current stimulation (Cecere *et al.*, 2015). Note however that our study was not specifically designed for the purpose of these analyses. Our results should thus be considered exploratory.

In our data, EEG power spectra during stimulation versus immediately before and after stimulation (peri-stimulation) showed comparable properties, characterised by a prominent alpha peak at around 10 Hz in all cases (*Figure 4a-d*). Topographical representations (averaged across attention conditions for data during stimulation) indicated that the alpha power distribution was highly similar and electrode clusters derived from alpha lateralisation analyses covered bilateral maxima. For further analyses, we determined individual alpha frequency (IAF) and power (IAP) from log-power spectra averaged across electrode clusters. We then tested whether peri-stimulation measures predicted online measures and whether predictions differed between stimulation conditions. Regarding peak frequency, regression analyses showed that peri-stimulation IAF was a significant predictor of IAF during theta-(test vs constant model: F(1,15) = 16.066, *P* = 0.00114), alpha-(F(1,15) = 20.285, *P* = 0.00042) and beta-band stimulation (F(1,15) = 15.494, *P* = 0.00132). This was further expressed in greater-than-zero regression slopes (*β*_*1*_) as listed in *Table 1* (Note that *Table 1* also lists regression intercepts (*β*_*0*_) for completeness).

**Table 1.** Parameters of robust linear regressions for all stimulation conditions. Predictor = alpha measure (IAF, IAP) peri-stimulation; outcome = alpha measure during stimulation. 95% confidence intervals (*CI*) were bootstrapped (*N* = 10000) using the bias-corrected-and-accelerated method.

| | *Stimulation* | $R^2$ | $\beta_0$ | 95% CI | $\beta_1$ | 95% CI |
| --- | --- | --- | --- | --- | --- | --- |
| IAF | Theta | 0.517 | -0.145 | [-0.420 0.357] | 0.615 | [0.197 0.900]* |
|  | Alpha | 0.575 | -0.011 | [-0.339 0.336] | 0.787 | [0.401 1.033]* |
|  | Beta | 0.508 | -0.107 | [-0.383 0.322] | 0.619 | [0.189 0.994]* |
| IAP | Theta | 0.792 | 1.968 | [-1.679 3.421] | 1.293 | [0.722 1.972]* |
|  | Alpha | 0.761 | 2.256 | [-3.481 3.502] | 1.338 | [0.434 1.752]* |
|  | Beta | 0.768 | 2.643 | [-0.774 3.862] | 1.461 | [0.749 2.032]* |

IAF – individual alpha frequency; IAP – individual alpha power; $R^2$ – goodness of fit; $\beta_0$ – intercept; $\beta_1$ – slope; \* denotes significance

**Figure 4.**
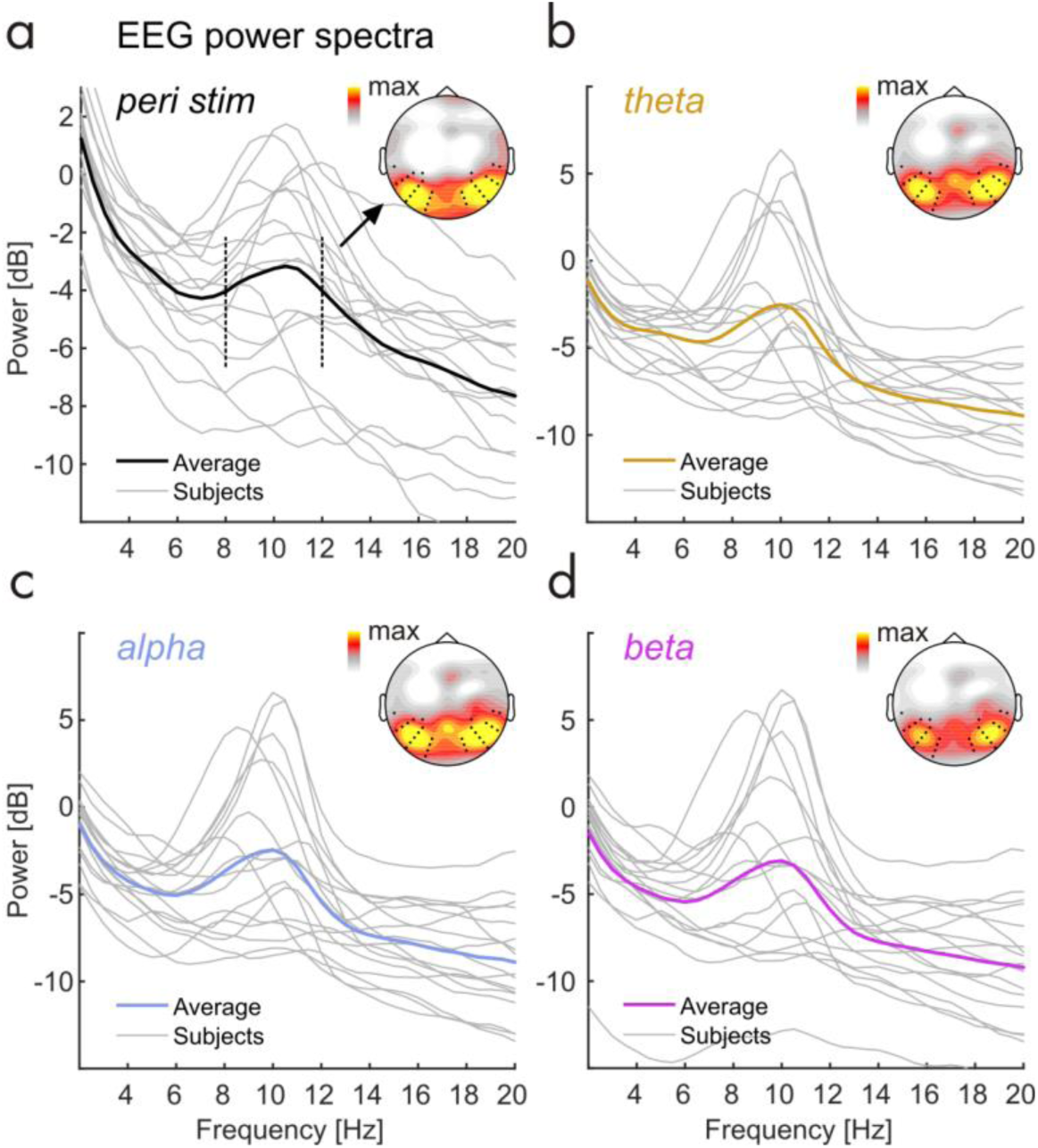
Individual EEG power spectra peri- and during stimulation. (**a**) Individual EEG power spectra (grey lines) peri-stimulation. Strong Black line shows group average. Inset scalp malp depicts topographical distribution of alpha power (8-12 Hz, frequency range indicated by black dashed vertical lines in spectrum). Black dots in scalp maps represent electrodes of the two clusters identified in alpha lateralisation analyses (see Figure 3). (**b**,**c**,**d**) Same as in (a) but during theta-(orange), alpha-(blue) and beta-stimulation (purple) respectively. Note that scales differ between (a) and (b),(c),(d).

Pairwise comparisons of regression slopes did not show any systematic differences in IAF linear dependencies between stimulation conditions (also see Figure 5a – c). More specifically, we did not find a significantly shallower regression slope for IAF during alpha-band than during theta-band stimulation (Δ*β*_*1*_ = −0.171, CI_BF_ = [-0.480 0.164]), or than during beta-band stimulation (Δ*β*_*1*_ = −0.003, CI_BF_ = [-0.467 0.161]). Linear dependencies were also comparable between theta- and beta-band stimulation (Δ*β*_*1*_ = 0.168, CI_BF_ = [-0.271 0.455]). Distributions of distances (absolute differences) of IAFs from the centre frequency of the alpha band stimulation (10.5 Hz) confirmed this finding (see *Figure 5d*): Individual distances were distributed equally across stimulation conditions (non-parametric Friedman test: X^2^(3,64) = 0.470; *P* = 0.926).

**Figure 5.**
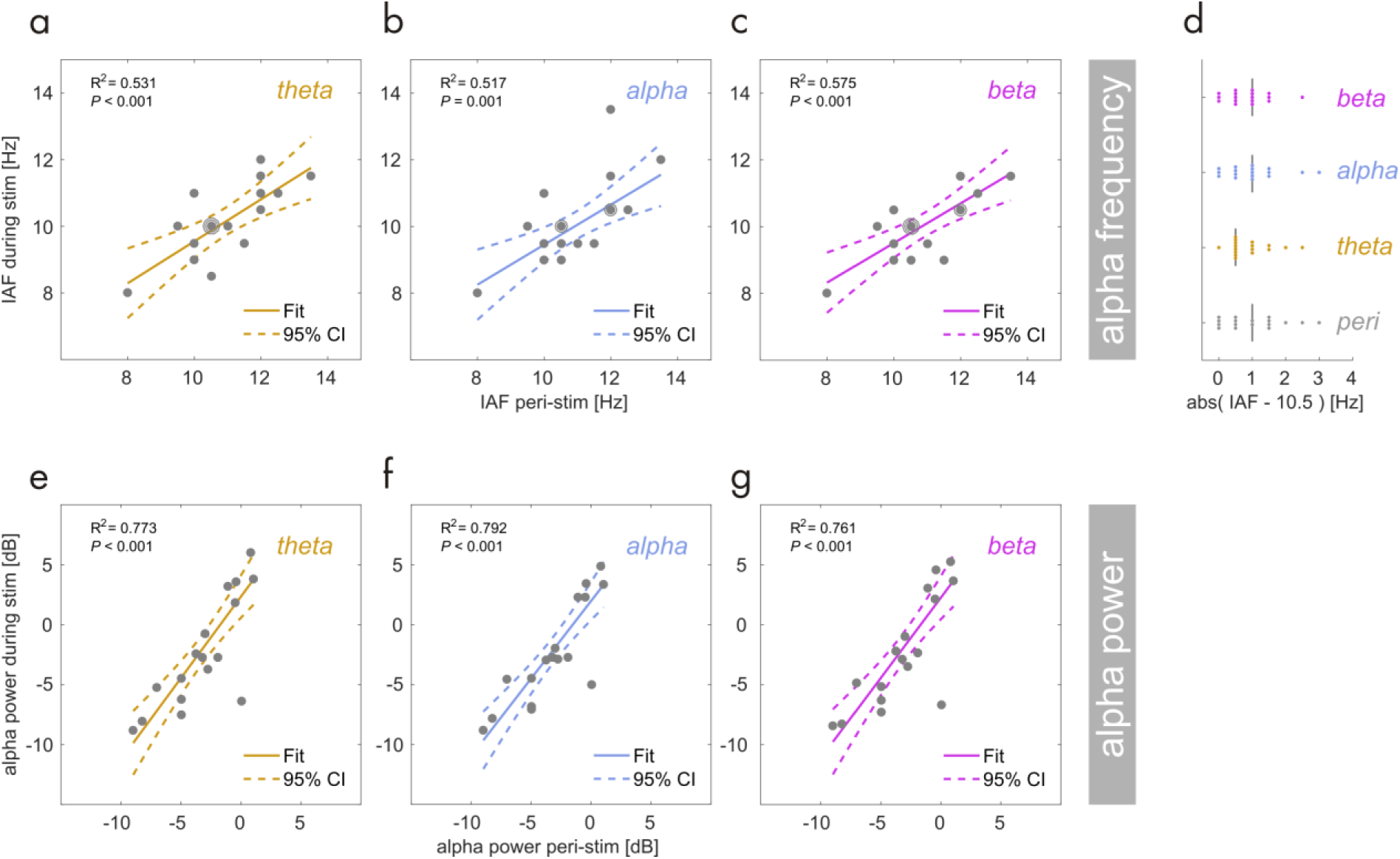
Linear regression: Individual alpha frequency (IAF) during theta-band (**a**), alpha-band (**b**) and beta-band (**c**) stimulation (y-axis) as a function of IAF peri-stimulation (x-axis). Grey dots represent participants. Dots with halos indicate more than one participant (max n = 3). Coloured lines depict a straight line fit and its confidence interval (dashed lines). Goodness-of-fit of the linear model provided as R^2^ along with corresponding *P*-Value. (**d**) Absolute distance of the individual alpha frequency from the centre frequency of the alpha band stimulation (10.5 Hz). Single coloured dots represent individual participants and vertical black lines show medians. Compare distances between alpha-band, other stimulation conditions and peri-stimulation. (**e-g**) Same as in (a) to (c) but for spectral power (in dB) at IAF.

Regarding power, peri-stimulation IAP systematically predicted IAP during stimulation for theta-(F(1,15) = 57.077, *P* < 0.0001), alpha-(F(1,15) = 47.628, *P* < 0.0001) and beta-band stimulation (F(1,15) = 49.677, *P* < 0.0001). *Figure 5e – g* depicts linear fits superimposed on individual data and displays corresponding Goodness-of-fit (R^2^). Regression slopes (*β*_*1*_) were all significant but intercepts (*β*_*0*_) were statistically indiscernible from zero (*Table 1*), thus contradicting a general increase in alpha power – indicative of alpha resonance – in any of the conditions.

### Behavioural performance not coupled to individual alpha power

In the original analysis of the behavioural data, reported in Keitel *et al.* (2017c), we found overall differences in performance between stimulation frequencies. Put briefly, participants detected target stimuli less accurately during beta-band stimulation and reaction times were found to be increased during theta-band stimulation.

Here, we concentrated on how individual alpha rhythms related to measures of performance as a function of stimulation frequency range. Note first that alpha lateralisation did not depend on alpha power in any of the three stimulation conditions (test vs constant model: maximum F(1,13) = 3.321, minimum *P* = 0.0915, *R*^*2*^ = 0.203, alpha band stimulation condition [two outlier removed by means of Cook’s distance]).

When looking at the relationship between the proportion of correct responses and alpha power per condition (*Figure 6a – c*), regression analyses revealed a systematic linear dependency during alpha-band stimulation (F(1,14) = 8.850, P = 0.010, *R*^*2*^ = 0.387, one outlier removed) but not during theta-(F(1,14) = 0.182, P = 0.679, *R*^*2*^ = 0.013, one outlier removed) and beta stimulation (F(1,14) = 0.369, P = 0.554, *R*^*2*^ = 0.026, one outlier removed).

**Figure 6.**
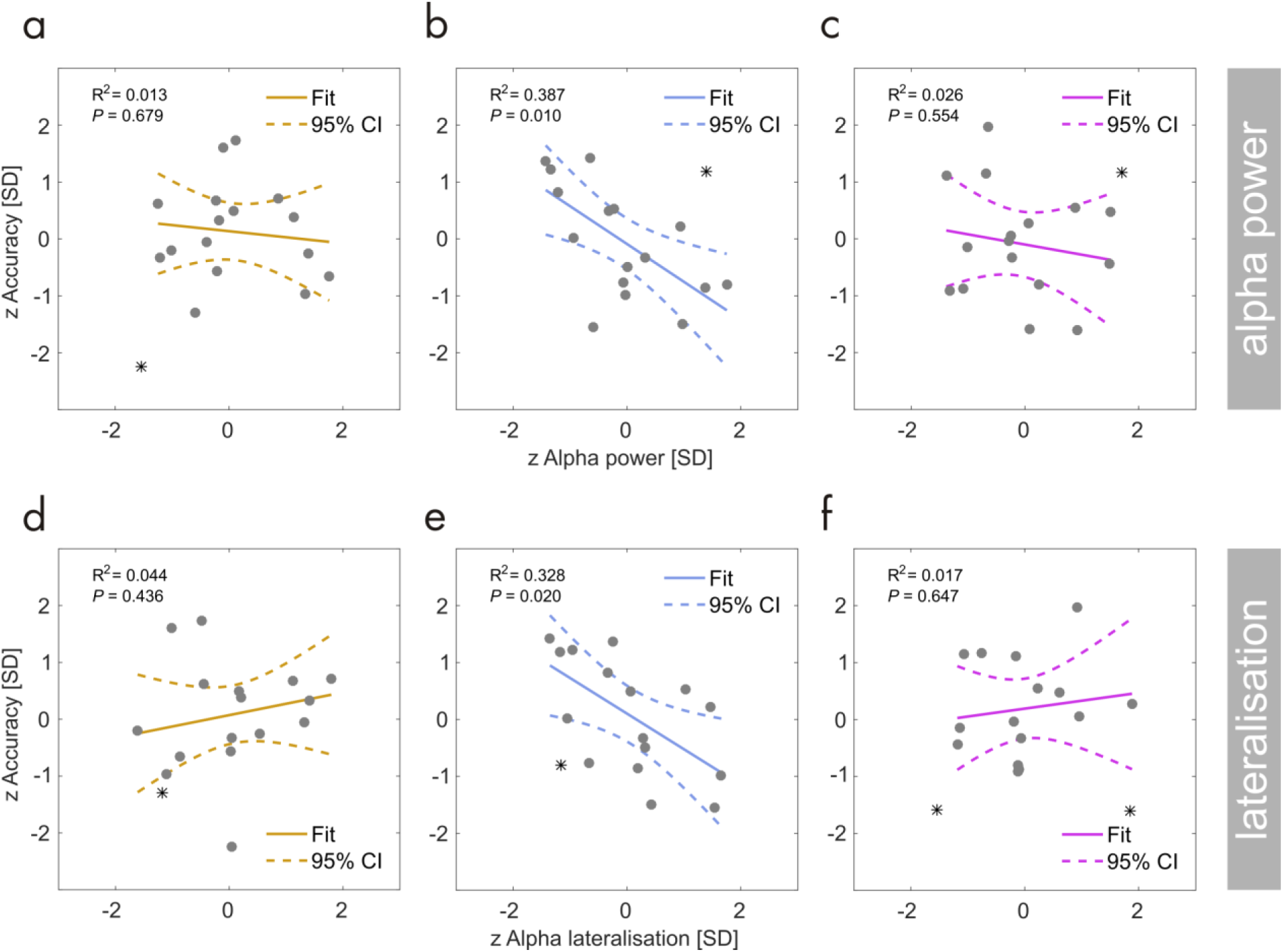
Linear regression: Individual performance in the behavioural task (z-scored, y-axis) during theta-band (**a**), alpha-band (**b**) and beta-band (**c**) stimulation as a function of total alpha power (z-scored, x-axis). Grey dots represent participants. Coloured lines depict a straight line fit and its confidence interval (dashed lines). Goodness-of-fit of the linear model provided as R^2^ along with corresponding *P*-Value. (**d-f**) Same as in (a) to (c) but for performance as a function of alpha power lateralisation (z-scored).

A closer examination of the significant linear fit during alpha band stimulation confirmed a significant negative regression slope (*β*_*1*_ = −0.664, 95% CI = [−1.226 -0.039]) meaning that participants with lower alpha power tended to perform better during alpha-band stimulation. Direct paired comparisons however revealed that regression slopes did not differ systematically between the stimulation conditions (theta vs alpha: Δ*β*_*1*_ = 0.588, *CI*_*BF*_ = [-0.809 1.606], two outlier removed; alpha vs beta: Δ*β*_*1*_ = −0.593, *CI*_*BF*_ = [-1.791 1.619], two outlier removed).

A similar overall pattern emerged from regressing correct-response proportions from alpha lateralisation (*Figure 6d-f*). We found a linear dependency during alpha-band stimulation (F(1,14) = 6.835, P = 0.020, *R*^*2*^ = 0.328, one outlier removed) but not during theta-(F(1,14) = 0.643, P = 0.436, *R*^*2*^ = 0.044, one outlier removed) and beta stimulation (F(1,13) = 0.230, P = 0.647, *R*^*2*^ = 0.017, two outlier removed). However, the regression slope for the alpha band stimulation condition failed to reach significance (*β*_*1*_ = −0.625, 95% CI = [-1.135 0.073]) thus rendering further direct comparisons unnecessary.

Neither alpha power nor lateralisation predicted response speed (median reaction times) in any of the stimulation conditions.

## DISCUSSION

The present re-analysis of previously published data (Keitel *et al.*, 2017c) examined the influence of quasi-rhythmic ongoing visual stimulation with frequencies in theta-(4 – 7Hz), alpha-(8 – 13Hz), and beta-bands (14 – 20Hz) on the EEG-recorded parieto-occipital alpha rhythm. We found that alpha power showed the typical effects of hemispheric lateralisation when participants attended to left or right stimuli irrespective of visual stimulation frequencies. Also, we compared individual alpha frequency and power measured immediately before and after stimulation (peri-stimulation) with alpha frequency and power during stimulation and failed to reveal effects consistent with a stimulus-driven entrainment of the intrinsic alpha rhythm (Thut *et al.*, 2011; Herrmann *et al.*, 2016a). Below we discuss the implications of our findings as well as possible reasons for why intrinsic alpha remained largely unaffected by our visual stimulation.

### Typical functional characteristics of alpha prevail during stimulation

Following the presentation of the cue on each trial, alpha power was modulated in a well-known fashion: A relative decrease in alpha power over parieto-occipital cortices contralateral to the cued location and an ipsilateral increase indicated corresponding covert shifts of spatial attention (Worden *et al.*, 2000; Rihs *et al.*, 2007; Capilla *et al.*, 2012) the quasi-rhythmic stimulation notwithstanding. Alpha power thus reflected the anticipatory attentional biasing of location-specific neural populations in early visual cortices (Kelly *et al.*, 2006; Thut *et al.*, 2006; Snyder & Foxe, 2010; Gould *et al.*, 2011). Higher alpha power ipsilateral to the attended position likely indicated suppression in visual cortices that received task-irrelevant input. Lower contra-lateral alpha power indexed a relative facilitation of visual input. In the present study, alpha thus retained its known functional characteristics during quasi-periodic visual stimulation irrespective of its spectral content.

Further, we found an imbalance in total alpha power between hemispheres. Higher right-than left-hemispheric alpha power has been consistently observed before (Slagter *et al.*, 2016; Benwell *et al.*, 2017a; Newman *et al.*, 2017) and has been linked to resource depletion within the right-hemispheric dorsal fronto-parietal attention network (the DAN; Sturm & Willmes, 2001; Corbetta & Shulman, 2011). Greater DAN activity relates inversely to alpha power (Sadaghiani *et al.*, 2012; Chang *et al.*, 2013; Zumer *et al.*, 2014). A depletion of DAN resources over the course of the experiment within the right hemisphere may have produced the here observed alpha power imbalance between hemispheres. Although an interesting effect, it was not in the focus of the current study and was thus not further investigated.

Previous studies have also demonstrated linear relationships between behavioural performance and alpha power, i.e. higher power coincides with lower hit rates (Hanslmayr *et al.*, 2005; van Dijk *et al.*, 2008), or its retinotopic lateralisation, i.e. greater lateralisation accompanies higher hit rates (Thut *et al.*, 2006), in target detection tasks. These dependencies were largely absent in our data, possibly due to the specific analysis approach employed here and our measure of behavioural performance. First, alpha power and lateralisation were evaluated on trials without target presentations making them only an indirect indicator of cortical processing at the time of target onset. Secondly, we measured behavioural performance as the proportion of correct responses, which is a measure of individual accuracy. However, alpha power variability, as an index of baseline cortical excitability (Romei *et al.*, 2008; Haegens *et al.*, 2014), may rather influence the response/perceptual bias, i.e. whether participants report detection or not regardless of the veracity of the percept (Limbach & Corballis, 2016; Benwell *et al.*, 2017a; Iemi *et al.*, 2017; Samaha *et al.*, 2017). Our analysis may thus have been insensitive to other alpha power and performance dependencies.

### No evidence for entrainment of the intrinsic visual alpha rhythm by quasi-periodic stimuli

We expected to observe two phenomena, alpha phase locking and alpha resonance if our stimuli were to entrain intrinsic (visual) alpha rhythms. Stimulation in the alpha frequency band indeed produced significant phase locking – but similarly so did theta and beta stimulation frequencies (see Keitel *et al.*, 2017c). Also, theta stimulation produced stronger phase-locking than alpha stimulation. This precise reflection of stimulus spectral content in the EEG, with a monotonous gradient of decreasing phase-locking from lower to higher frequency bands, speaks against a special proneness of alpha generators in visual cortex (Rosanova *et al.*, 2009; Herring *et al.*, 2015; Keitel & Gross, 2016) to entrain to visual quasi-periodic alpha band stimulation.

Alpha resonance, in turn, should have expressed as a greater increase in alpha power during stimulation within the IAF range (Fedotchev *et al.*, 1990; Schwab *et al.*, 2006) than theta and beta power during theta- and beta-band stimulation, respectively. Entrainment models explain such non-linear response gains through a population-wide phase alignment, whereby neuronal oscillators with a preferred frequency in the alpha band synchronize with the external drive. This mass synchronization would then shape a rhythmic neural response that can be measured macroscopically (Thut *et al.*, 2006). Evidently, our stimulation failed to produce a detectable alpha resonance. Alpha power was not significantly increased during alpha-band stimulation as compared with theta- or beta-band stimulation. This null-effect further translated into comparable alpha power lateralisation during sustained attention. Additionally, “baseline” alpha power in the absence of stimulation predicted individual alpha power during stimulation similarly in all three quasi-rhythmic conditions. Bayesian inference confirmed that an effect of stimulation frequency on alpha power was implausible. Note however that demonstrating entrainment effects on power statistically requires higher signal-to-noise ratios than other spectral measures, such as inter-trial phase consistency (Ding & Simon, 2013).

The absence of alpha resonance effects may have been a direct consequence of a lack of phase locking of intrinsic alpha generators: If intrinsic alpha (as opposed to stimulus-related neural activity) entrained to the external drive, this should have led to a synchronization of intrinsic alpha to stimulation. Entrainment should therefore entail an overall shift in individual alpha frequencies (IAFs) when these do not match the spectral profile of the stimulation. However, alpha band stimulation did not pull IAFs towards the stimulation centre frequency (10.5 Hz) in our case. Instead, IAF measured in the absence of stimulation predicted IAF during stimulation comparably well, irrespective of stimulation frequency.

Note however that these effects on frequency may be small and escape the here achieved spectral resolution (0.5 Hz). They may also depend non-linearly on the difference between IAF and stimulation centre frequency, such that participants with IAFs further removed from the centre frequency (10.5 Hz) may be less prone to experience a pull effect. Further, considering that our stimulation was only quasi-rhythmic, i.e. occurred within a band around but not at the centre frequency may have weakened a pull effect additionally. Lastly, work on the role of intrinsic rhythms in tactile perception has led to the argument that focussing on individual peak frequencies only may neglect effects on rhythmic activity beyond the peak (Baumgarten *et al.*, 2017). Also note that the EEG is likely dominated by a strong parieto-occipital alpha generator. However recent research suggests more than one concurrent alpha rhythmic ‘mode’ involved in visual processing (Keitel & Gross, 2016; Barzegaran *et al.*, 2017). It is therefore possible that our stimulation failed to influence the dominant rhythm but affected a less prominent alpha component that is more difficult to observe given the sparse spatial resolution of EEG recordings. Dedicated source reconstructions in future MEG experiments may shed further light on this issue.

### Current results suggest boundary conditions for the entrainment of endogenous alpha generators in the visual system

Taken together, we did not find that intrinsic alpha rhythms entrained to our low intensity quasi-rhythmic visual stimuli at alpha frequency. It is not clear whether this was a direct consequence of the quasi-rhythmicity of our stimulation. If so, our results would be consistent with a recent claim that only strictly periodic stimulation can produce alpha entrainment (Notbohm & Herrmann, 2016). Alternatively, stimulation intensity could have been too low to force cortical alpha generators to follow the entrainment regime (Pikovsky *et al.*, 2003; Thut *et al.*, 2011; Herrmann *et al.*, 2016a). In line with this assumption, a recent study delivered separate rhythmic trains at rates around (and including) individual alpha frequencies with an orthogonal luminance manipulation and demonstrated the positive influence of stimulus intensity on entrainment (Notbohm *et al.*, 2016). Note that in Notbohm *et al.* (2016), for stimulus intensities greater than 300 cd/m^2^ entrainment occurred within a range of stimulus frequencies of IAF ± 2 Hz approximating the dynamic bandwidth of our alpha-band stimulation (8-13 Hz). The intensity however exceeded the maximum luminance of the stimulation employed here (∼30 cd/m^2^) by one order of magnitude. Put differently, our stimulation intensity would have had to be increased ten-fold to expect entrainment effects to emerge while assuming that variations in frequency do not influence the process of entraining endogenous alpha.

Studies reporting neural phenomena consistent with alpha entrainment typically used stimulation parameters that feature one or more of the following properties: Stimulation was of high intensity and employed strictly rhythmic on-off flicker. Also, entraining stimuli were typically presented centrally and therefore close to the fovea (Notbohm & Herrmann, 2016; but see Sokoliuk & VanRullen, 2016) and in isolation (Mathewson *et al.*, 2012; but see Gulbinaite *et al.*, 2017), i.e. in the absence of potentially competing stimuli (Keitel *et al.*, 2013). In contrast, our stimulation consisted of two para-foveal stimuli presented laterally at low intensity. Although luminance was modulated locally, it was the global stimulus contrast that varied (quasi-) sinusoidally over time. In an attempt to formulate boundary conditions, we thus suggest that low-intensity, non-foveal and frequency-varying stimulation, as employed here, may be insufficient to produce purely stimulus-driven entrainment of endogenous alpha rhythms in the visual system.

Physical stimulus properties aside, previous research has also argued for a top-down aspect in entrainment: rhythmic stimulus presentations may not only entrain brain oscillations directly (bottom-up) but generate predictions about future stimulus occurrences (Nobre, 2001; Thut *et al.*, 2011; Wiener & Kanai, 2016; Breska & Deouell, 2017). In that case, entrainment may be realised through recurring feedback loops that connect higher-order with early visual cortices (Samaha *et al.*, 2015). A candidate network providing this feedback might be the dorsal fronto-parietal attention system (Corbetta & Shulman, 2011). The quasi-rhythmic nature of our stimulation may have therefore prevented strong and precise temporal predictions about the temporal evolution of the stimulus. Also, the randomness in stimulus frequency fluctuations likely impeded memory-based predictions (Breska & Deouell, 2017; also see Obleser *et al.*, 2017). Moreover, the waxing and waning of stimulus contrast in our design was of little relevance for performing in the behavioural task. Stimulation phase had no predictive value for target presentation, and target duration (0.3 sec) exceeded the length of a 4-Hz cycle (.25 sec, i.e. the lower boundary of the lowest stimulation frequency used in this study), further precluding potential entrainment effects.

These collective arguments for the absence of intrinsic alpha entrainment effects in our study notwithstanding, we emphasise that we found substantial EEG phase-locking to the stimulation. This neural response was highly specific to the spectral composition of each stimulus (Keitel *et al.*, 2017c) and demonstrated that stimulation was generally strong enough to elicit neural responses. Weighing the absence of alpha entrainment effects against the substantial EEG-stimulus locking in our study, we suggest that the presence of a frequency-following response to a stimulus alone is not sufficient to conclude that intrinsic rhythms, specifically alpha, have been entrained by the stimulation (Keitel *et al.*, 2014).

### Boundary conditions challenge role of stimulus-driven visual alpha entrainment in sensory sampling

The current concept of the role of entrained intrinsic rhythms in perception largely borrows from research into auditory perception. Rhythmic, and thus deterministic, auditory stimuli afford efficient neural encoding (Large & Jones, 1999; Schroeder *et al.*, 2010; Henry & Herrmann, 2014). For these type of stimuli, entrainment enables precise predictions of future stimulus occurrences through low-frequency brain-stimulus synchronisation (Henry & Obleser, 2012; Henry *et al.*, 2014). Similar effects of rhythmic stimulation have since been reported for visual perception (Cravo *et al.*, 2013; Calderone *et al.*, 2014; Spaak *et al.*, 2014; Sokoliuk & VanRullen, 2016).

Entrainment likely plays a fundamental role in speech comprehension (Giraud & Poeppel, 2012). Components of the auditory speech signal, such as prosody, phoneme and syllabic rate, entrain low-frequency rhythmic activity in auditory cortices (Gross *et al.*, 2013; Peelle *et al.*, 2013; Keitel *et al.*, 2017a; Keitel *et al.*, 2018). The strength of entrainment thereby influences comprehension performance (Zion Golumbic *et al.*, 2013; Rimmele *et al.*, 2015; Zoefel & VanRullen, 2015). More recent studies have found that visual components of speech, such as lip movements (Chandrasekaran *et al.*, 2009) and gestures (Biau & Soto-Faraco, 2015), produce similar entrainment effects in visual cortices (O’Sullivan *et al.*, 2016; Park *et al.*, 2016) that serve to enhance auditory signals under challenging listening conditions (Zion Golumbic *et al.*, 2013).

Most important in the present context: speech is naturally quasi-rhythmic. This vital property allows it to convey information. Apart from speech, quasi-rhythms typically compose natural dynamic visual scenes (Kayser *et al.*, 2003; Butts *et al.*, 2007; Mazzoni *et al.*, 2011) and it has been argued that they benefit visual perception more than strictly rhythmic input (Buracas *et al.*, 1998; Blake & Lee, 2005). Therefore, in an entrainment framework of visual perception it would seem sensible to assume that the cortical generators underlying intrinsic rhythms should resonate with a range of stimulus frequencies even under low-intensity conditions. Consequently, this would allow them to accommodate dynamic changes in quasi-rhythmic input within their preferred frequency range, at least to some extent. It is thus disputable whether visual alpha entrainment can fulfil a role similar to low-frequency entrainment in speech perception, when it can only be produced by strictly periodic stimulation under certain conditions (high intensity, foveal presentation) in an experimental setting. We point out that drawing immediate parallels between how auditory and visual systems exhibit entrainment entails the implicit assumption that both respond similarly to rhythmicity in sensory input. Challenging this notion, Zoefel and VanRullen (2017) have recently argued that vision and audition may rely on distinct internal sampling strategies that exploit stimulus rhythmicity differently. In contrast to the auditory system, phase-locking to a dynamic stimulus may not be an integral part of the attentional selection process in vision. Instead, the visual system could maintain an autonomous visual sampling strategy that is independent of the stimulus and whose neural signature is the parieto-occipital alpha rhythm. Our paradigm likely promotes such a strategy because the stimulus dynamics are behaviourally irrelevant. Also, we found a tendency of EEG phase-locking to theta-band stimulation to be reduced on trials in which alpha power was high (see *Supplementary Material*) further suggesting an interplay in cortical visual processing that may rely on the autonomy of stimulus-driven and intrinsic rhythms. In line with this autonomy assumption, van Wassenhove (2016) has recently proposed that the ability of intrinsic rhythms to decouple from (periodicities in) external stimulation may be a vital prerequisite for a stimulus-independent monitoring of the passage of time.

In summary, our findings do not support an entrainment of the intrinsic parieto-occipital alpha rhythm driven by quasi-periodic visual stimulation in the alpha band. We have discussed a number of mitigating factors (quasi-periodicity, low-intensity, task irrelevance of the stimulation) that may have contributed to the absence of effects. We suggest that future studies into visual entrainment may need to take into account that phase synchronisation of neural activity with visual stimulation per se is a necessary but insufficient condition to infer entrainment of intrinsic brain rhythms.

## ACKNOWLEDGMENTS

Funded by a Wellcome Trust Senior Investigator Grant awarded to GT and JG (#098433/#098434). Lucy Dewhurst and Jennifer McAllister assisted in data collection. We thank Anne Keitel for comments on the manuscript. The experimental stimulation was realized using Cogent Graphics developed by John Romaya at the Laboratory of Neurobiology, Wellcome Department of Imaging Neuroscience, University College London (UCL).

## Supporting Information

Additional supporting information can be found in the online version of this article.

## Competing interests

The authors declare no competing interests.

## Author contributions

CK designed research, performed research, analysed data and wrote the article. JG designed research analysed data and wrote the article. CSYB and GT designed research and wrote the article.

## Data accessibility

EEG data are available on the Open Science Framework (OSF) under the URL osf.io/apsyf (Keitel *et al.*, 2017b)

## Footnotes

1 Note that, entrainment’ and, phase synchronisation’ are frequently used synonymously in the literature. To avoid confusion, we will use, phase-locking’ or, phase synchronisation’ whenever we refer to neural responses that are largely stimulus-driven and, entrainment’ when it is assumed that phase-aligned endogenous rhythms contribute to the measured neural response.

2 These 17 were selected from a total of 22 recorded participants. Exclusion criteria are reported in the original study.

