## Supplementary Materials for "No changes in parieto-occipital alpha during neural phase locking to visual quasi-periodic theta-, alpha-, and beta-band stimulation"

*Estimating individual alpha frequencies (IAF)*

IAF's were estimated from EEG power spectra as shown in *Figure 4*. We found that taking the second order gradient of raw power spectra and subsequently applying a Savitzky-Golay Filter (Savitzky & Golay, 1964), based on a third-order polynomial and with a frame length of 9 consecutive frequency bins, allowed for maximum peak identification within the frequency range 6 – 16 Hz for each participant (see *Figure S1*). In MATLAB this was implemented as:

```
smSpec = sgolayfilt(-gradient(gradient(powSpec)),polyOrder,filterLength);
```

taking `powSpec`, i.e. a numerical vector containing spectral power estimates for frequencies in ascending order, as input. Inputs to the `sgolayfilt` function were the order of the polynomial filter kernel (`polyOrder`) and its length (`filterLength`). We used the default setting of '3' for the order of the polynomial. The filter length was chosen based on our spectral resolution (0.5 Hz) meaning that each point-estimate depended on the neighbouring ±4 frequency bins (±2 Hz).

A similar step has been implemented in a recent algorithmic and more comprehensive approach of IAF determination (Corcoran *et al.*, 2017).

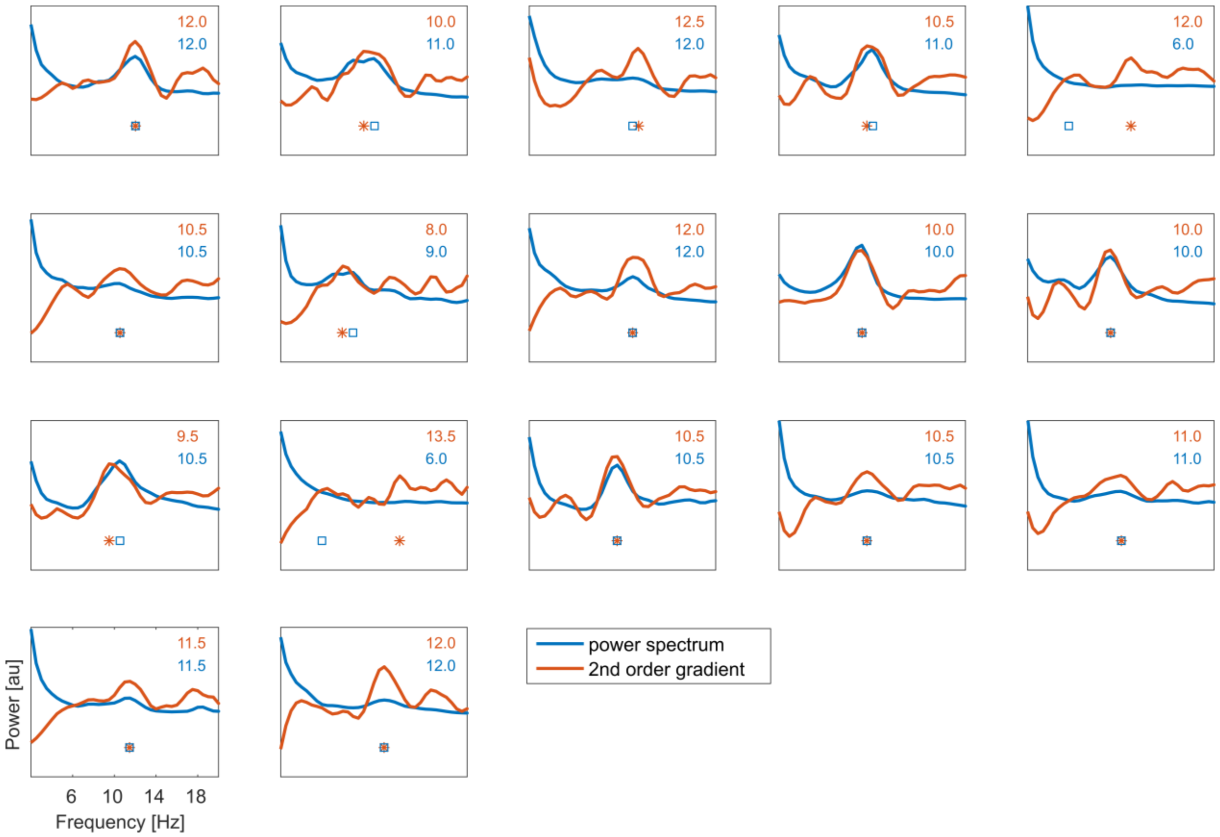


**Figure S1** Identifying individual peak alpha frequencies (IAF). Each plot shows the EEG power spectrum (blue line) collapsed across two electrode clusters covering parieto-occipital areas (see Figure 3b) for one participant in the absence of ('peri') visual stimulation. Most of them show clear peaks at frequencies within the alpha band (blue numbers, top right corner) that can easily be identified (also see blue squares below spectra). For these participants, power spectrum IAF largely coincides with an estimate (red numbers) based on the Savitzky-Golay-filtered second-order gradient of the power spectrum (red line). For a few participants however, only the latter provides a sensible estimate of the IAF. For IAF analyses we thus opted for IAF estimates based on the gradient spectra (red lines, numbers and asterisks).

*Single-trial variability of alpha lateralisation and EEG-stimulus locking*

In an additional analysis we turned to how alpha activity influenced sensory processing of continuous visual input. According to the inhibitory role ascribed to alpha activity (Jensen & Mazaheri, 2010), sensory processing as measured by EEG-stimulus locking should normally be attenuated during periods of relatively high alpha power (*Figure S2a & b*). This inverse relationship could be modulated by an external alpha pacemaker, such as our alpha band stimulation, interacting with the intrinsic rhythm.

For this purpose, 1-sec epochs of EEG data during stimulation underwent processing and spectral decomposition as described in Keitel *et al.* (2017). Alongside single-trial alpha power estimates, analyses yielded a measure of EEG-stimulus locking indexing early cortical visual processing. This EEG-stimulus locking was quantified as phase cross-coherence (XCOH) between stimuli and brain responses. XCOH specifically indicated the consistency of stimulus-brain phase differences as a statistic across trials (Cohen, 2014; Gross, 2014).

Enabling an examination of single-trial variability of XCOH thus required applying a methodological approach proposed by (Richter *et al.*, 2015): Quantities that can only be calculated and meaningfully interpreted across trials (such as XCOH) normally prevent single-trial analyses. For the purpose of correlating it with another measure that can be extracted from single trials (alpha power) a single-trial XCOH surrogate can be extracted by means of a Jackknife procedure. Hereby, XCOH is estimated *n* times (with *n* being the number of trials) across *n*-1 trials of one condition successively leaving out each trial once. Across Jackknife estimates of XCOH a variability is retained that can be explained by the contribution of each (left-out) trial to the original XCOH estimate based on all *n* trials. For our case, this property enabled an investigation of dependencies between within-subject variations in alpha power and XCOH.

To sample both measures in a comparable manner, similar to alpha power, XCOH Jackknife estimates were pooled over sensors within lateralised occipito-parietal electrode clusters contralateral to the driving stimulus (see *Figure 3b*). More specifically, the left cluster of electrodes sampled XCOH with the right stimulus and vice versa. Note that this spatial sampling of XCOH likely attenuated effects when analysing the beta-band condition because only few sensors within these clusters showed significant XCOH with beta-band stimulation.

We further collapsed alpha power over frequencies IAF ± 1 Hz and applied the same bandwidth when aggregating XCOH around respective centre frequencies (5.5 Hz for theta-, 10.5 Hz for alpha-and 17 Hz for beta-stimulation conditions).

Following these steps, estimates of single trial variability of alpha power and XCOH ipsi- and contralateral to the respective attended position were obtained. As a control, we tested whether XCOH averaged across Jackknife estimates showed a pattern of results consistent with our previous report in which we employed a different analysis approach (Keitel *et al.*, 2017). Average XCOH was subjected to a three-way repeated-measures ANOVA with factors of *hemisphere* (contra- vs ipsilateral), *stimulus frequency* (theta, alpha, beta) and *stimulus position* (attend left vs right). As laid out in the Results section below, this ANOVA confirmed our earlier findings.

Finally, for each participant and condition, single trial alpha power was used to predict EEG-stimulus locking (XCOH). For that purpose, we fitted robust linear models, as described above, based on z-scored alpha power (dB-scaled) and XCOH (log) for sensor clusters contra- and ipsilateral to each stimulus, experimental condition and participant separately. Linear fits yielded regression slopes that were entered into second-level statistics. We repeated this process with surrogate XCOHs calculated between EEG and time-reversed stimulus functions on each trial to ascertain that any effects of alpha power on XCOH were specific to stimulus-evoked responses.

A four-way repeated-measures ANOVA tested for influences of factors of *hemisphere* (contra- vs ipsilateral), *stimulus frequency* (theta, alpha, beta), *stimulus position* (attend left vs right) and *stimulus direction* (original vs reverse) on putative dependencies between alpha power and EEG-stimulus locking variability. Note that regression slopes were sign-flipped before the ANOVA because extracting only one (XCOH) but not the other covariate (alpha power) by means of Jackknife estimates reverses their sign (Richter *et al.*, 2015).

As documented below, the ANOVA only indicated an effect of *stimulus frequency*, which further interacted with *stimulus direction*. Data were thus collapsed across factors *hemisphere* and *stimulus position*. T-tests against zero (employing a Holm-Bonferroni correction of *P*-values) were conducted to establish whether the resulting slopes deviated significantly from zero. Further post-hoc t-tests were used for subsequent pairwise comparisons.

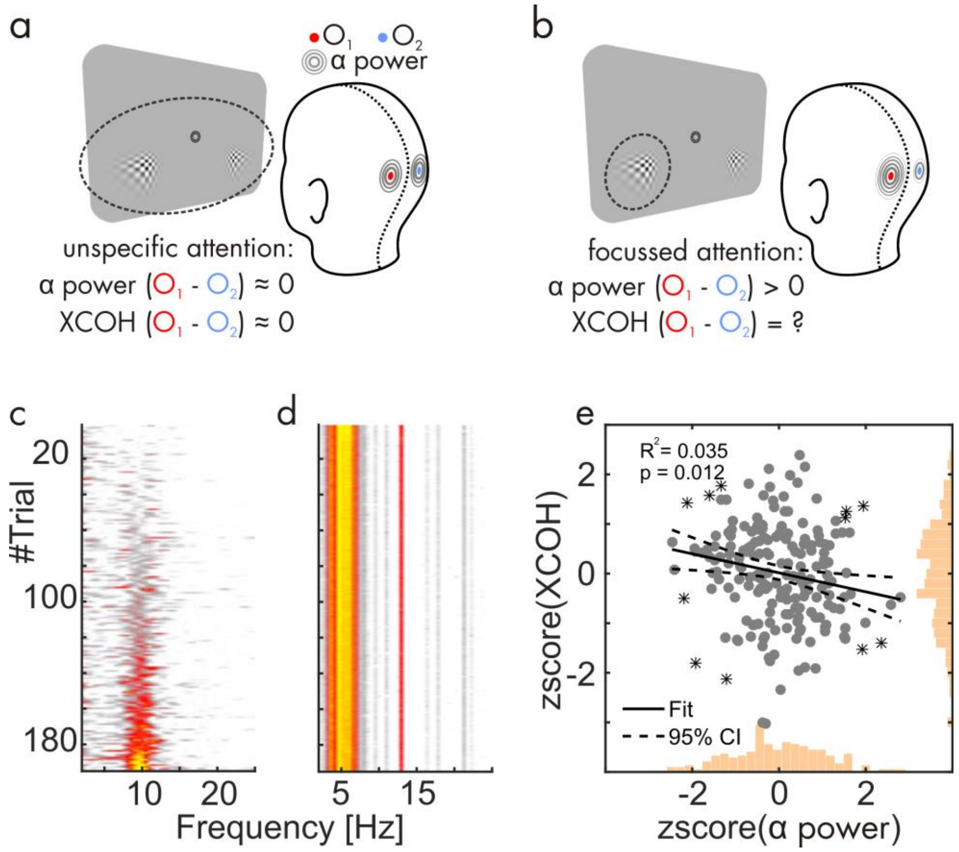


**Figure S2** Regressing cross-coherence (XCOH) from alpha power (**a**) Rationale – When attention is not specifically allocated to the left or right stimulus, alpha power and XCOH measured at retinotopically corresponding EEG sensors (schematically: $O_1$ and $O_2$) should be close to level. (**b**) When attention focuses on one stimulus position (dashed circle, left stimulus) alpha power is known to lateralise by increasing at ipsilateral

($O_1$) and decreasing at contralateral sites ($O_2$). Our regression analysis investigates whether single-trial variations in thus lateralised alpha power influence sensory processing measured as EEG-stimulus locking (XCOH). (**c**) Exemplary stacked spectra of one representative subject during visual theta-band stimulation and attending the right stimulus. Trials (y-axis) have been sorted according to power within the individual alpha frequency band (IAF +/- 1Hz) for display only. (**d**) Same subject and condition – XCOH spectra, stacked by trial and sorted according to XCOH in the 4 – 7 Hz band for display. The gradient from high to low lateralisation trials can hardly be seen because the applied Jackknife procedure greatly reduces variance in the data. (**e**) Scatterplot of z-scored data from (c) and (d). Each dot represents one trial. Asterisks denote outliers identified by means of Cook's distance. Straight black line depicts linear fit (dashed lines are 95% confidence intervals). Histograms on x- and y-axes depict approximately normal marginal distributions of alpha power and XCOH across trials.

*Alpha power predicts EEG-stimulus locking depending on stimulation frequency*

This analysis targeted possible influences of alpha power on neural phase locking (XCOH) to left and right stimuli (*Figure S2a – b*) as an index of sensory processing. Qualifying our approach, we found a pattern of results similar to the original study: XCOH decreased with increasing *stimulus frequency* ($F(2,32) = 52.63$, $P_{GG} < 0.001$, εGG = 0.996, $\omega_p^2 = 0.747$). Additionally, XCOH showed a trend to differ between recording sites at positions contra-vs ipsilateral to the driving stimulus ($F(1,16) = 4.07$, $P = 0.061$, $\omega_p^2 = 0.146$) with greater XCOH measured at sensors covering the hemisphere contralateral to the stimulation (*Figure S3a*). This was likely due to the spatial specificity, i.e. retinotopy, in early visual processing, that was not explicitly tested in the original study: a stimulus presented to the left hemifield is predominantly processed by the right hemispheric visual cortex and vice versa. XCOH magnitude was independent of whether the left or right stimulus was attended ($F(1,16) < 1$). Neither interaction term reached statistical significance ($F_{max}(2,32) = 2.13$, $P_{min} = 0.135$).

Alpha power showed variability across trials of respective conditions under conditions of focused spatial attention (illustrated for one representative participant during theta-band stimulation while attending the right stimulus in *Figure S2c*) as did XCOH (*Figure S2d*). Both alpha power and XCOH were normally distributed. This enabled linear regression analyses as exemplified in *Figure S2e*: Alpha power was used as a predictor for XCOH across trials. The solid black line represents a robust linear fit, the slope of which was entered into second-level statistical analyses.

*Figure S3a* depicts individual regression slopes schematically and grouped by stimulation condition and whether the respective XCOH driving stimuli were attended. Slopes averaged across participants are superimposed in colour.

Slope analyses revealed that linear relationships of XCOH and alpha power differed between *stimulus frequencies* ($F(2,32) = 9.24$, $P_{GG} = 0.006$, $\varepsilon_{GG} = 0.984$, $\omega_p^2 = 0.320$; also see *Figure S3b*). Slopes did not differ between contra- and ipsilateral *hemispheres, stimulus positions* or *stimulus direction* (all

F(1,16) < 1) but *stimulus direction* influenced the alpha-XCOH relationship specifically for some *stimulus frequencies* ($F(2,32) = 3.73$, $P_{GG} = 0.036$, $\varepsilon_{GG} = 0.977$, $\omega_p^2 = 0.135$; also see *Figure S3b)*. Other interactions remained insignificant ($F_{max}(1,16) = 2.69$, $P_{min} = 0.121$ for *stimulus direction* * *hemisphere*).

Contrasts of slopes, averaged across *hemisphere* and *stimulus position*, against zero (two-tailed t-tests) indicated that only theta-band phase locking with the originally seen (but not with the surrogate, reversed stimulus) systematically *decreased* with increasing alpha power (mean slope *M* = -0.032, standard error of the mean *SEM* = 0.011, t(16) = -3.15, $P_{HB} < .05$). All other slopes did not differ significantly from zero ($P_{HB} > .1$). In a direct comparison, slopes based on phase locking with original vs reversed theta band stimulation only differed marginally (t(16) = -1.44, *P* = .085, one-tailed, uncorrected) although slopes differed substantially for original theta vs alpha (t(16) = -3.91, *P* = .001, two-tailed, uncorrected) and theta vs beta stimulation (t(16) = -3.40, *P* = .004, two-tailed, uncorrected). Note again that the approach employed here did not sample XCOH to beta-band stimulation optimally thus potentially underestimating effects in this condition.

In sum, our analysis provides little evidence of a hypothesised inverse linear relationship between alpha power and EEG-stimulus locking. If at all, neural phase-locking to quasi-rhythmic stimulation in the theta band (4 – 7 Hz) decreased as a function of alpha power across trials. Interestingly, a similar relationship between sub-alpha phase locking to stimulation and alpha power has previously been reported in the auditory domain (Kerlin *et al.*, 2010; Wöstmann *et al.*, 2016). However, there was no clear statistical distinction between slopes based on original vs reversed theta-band stimulation. It is thus possible that our analysis is more sensitive to a general inverse relationship between alpha power and theta phase (locking) that is stimulus un-specific (Klimesch *et al.*, 1997; Lundqvist *et al.*, 2011).

There was no indication of a similar relationship between alpha power and phase locking to alpha stimulation. We speculate that this points towards (1) an interaction between intrinsic and stimulus driven rhythms, consistent with the notion of entrainment, that counteracts the expected inverse relationship, or (2) the fact that our analyses cannot fully separate stimulus-driven from intrinsic rhythmic activity when both fall within the same frequency range, which may have a similar effect.

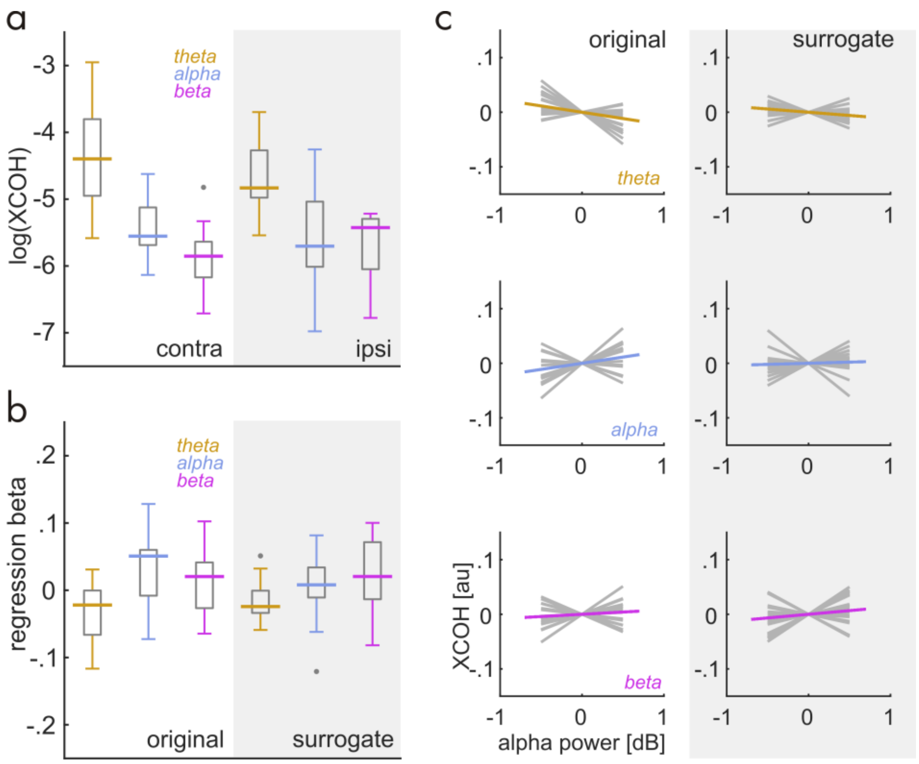


**Figure S3** Single-trial regressions of EEG-stimulus locking (XCOH) from alpha power. (**a**) Jackknife-derived XCOH estimates show a pattern similar to Keitel et al. (2017) – XCOH decreases with stimulation frequency (here: contra- as well as ipsilaterally to the stimulus position). (**b**) Distributions of regression coefficients (betas) for regression analyses based on XCOH computations using the original (left) and the time-reversed stimuli (right, "surrogate"). (**c**) Individual regression slopes (grey line bundles) split by theta-, alpha- and beta-band (top to bottom) stimulation and whether based on XCOH computations using the original (left) and the time-reversed stimuli (right, "surrogate"). Each line is the result of the analysis exemplified in Figure S2.

*Analyses including the strictly-rhythmic stimulation condition*

Data from the conditions with strictly rhythmic stimulation (left stimulus = 10 Hz, right = 12 Hz) were excluded from the analyses reported in the main text. The reason for the exclusion was that the constant-frequency conditions differ physically in more than one way from the frequency-varying conditions: no frequency changes over time (strictly rhythmic and thus single-frequency), different frequencies left and right. Here we present a parallel analysis including the strictly rhythmic stimulation condition and additional illustrations. Because of the limitations given above these results should be interpreted with caution.

A first glance at an overlay of the power spectra of all stimulation conditions illustrates that the frequency composition of the ongoing EEG during constant-frequency stimulation is largely comparable with the other conditions (*Figure S4a*). Focussing on the effects of spatial attention on alpha power yielded the typical retinotopic lateralisation (scalp map in *Figure S4b*). Also during constant-frequency stimulation alpha was significantly suppressed when participants attended to respective contralateral visual hemifields. Further note that total alpha power showed a topographical and spectral distribution similar to the other stimulation conditions as well as during peri-stimulation intervals (*Figure S4c*). We repeated the repeated-measures ANOVA (Bayesian version) on alpha power with factors of *hemisphere* (left vs right), attended *stimulus position* (ipsi- vs contralateral) and *stimulation frequency* (theta, alpha, beta & now including the constant conditions).

Again, Bayesian inference showed strong evidence for an influence of *attended position* on alpha power ($BF_{10}$ = 7.499 x $10^8$ ± 0.54%). This analysis also confirmed the strong evidence for an imbalance in overall alpha power between hemispheres ($BF_{10}$ = 1760.810 ± 0.46%). Crucially, the factor *stimulus frequency* showed substantial evidence in favour of $H_0$, i.e. no effect of stimulation frequency on alpha power ($1/BF_{10} = BF_{01}$ = 12.658 ± 0.37%). Comparing models, a linear combination of the factors *attended position* and *hemisphere* was most plausible given our data ($BF_{10}$ = 9.928 x $10^{12}$ ± 3.13%) and outperformed a linear combination additionally including the factor stimulus frequency by a factor of ~8 ($BF_{10}$ = 1.235 x $10^{12}$ ± 1.93%; ratio of model BFs = 8.036 ± 3.677%). In essence, these results confirm all findings from the analysis excluding the constant-frequency stimulation and therefore suggests that this type of stimulation does not have a specific influence on alpha power and its attentional modulation.

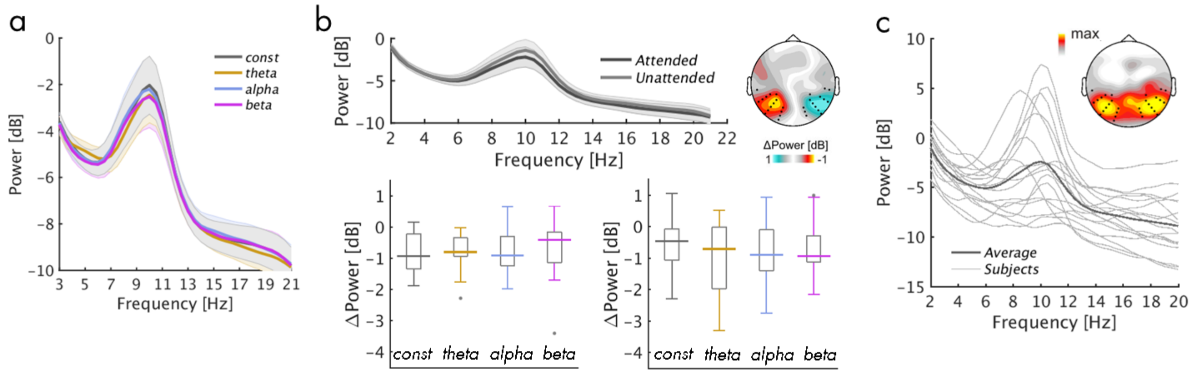


**Figure S4** EEG power spectra and effects of attention during constant-frequency stimulation (colour-coded grey in all plots). (**a**) EEG power spectrum superimposed with power spectra from frequency-varying stimulation conditions – compare with *Figure 1d*, rightmost panel. (**b**) Attention effects on alpha power. Top spectrum shows retinotopic alpha suppression with attention collapsed across hemispheres. Scalp map illustrates alpha power lateralisation. Box plots illustrate alpha suppression effects (Attended minus Unattended) in all four stimulation conditions for left and right electrode clusters (see scalp map), respectively. Contrasts against zero (two-tailed t-tests) all $P_{HB} < 0.05$ – compare with *Figure 3*. (**c**) Individual alpha power spectra collapsed across electrode clusters – compare with *Figure 4*.

Finally, we also conducted an analysis focusing on the relationship between alpha power and frequency during constant-frequency versus peri-stimulation. In the original analyses based on frequency-varying stimulation, we postulated effects specific to the alpha-band stimulation and relative to the other two stimulation conditions assuming entrainment. This analysis had to be adapted for the constant-frequency case because the two stimuli flicker at different frequencies (10 Hz left and 12 Hz right). We ran separate linear regressions of individual alpha power (IAP) and peak frequency (IAF) during stimulation from the same measures peri-stimulation for right- and left-hemispheric electrode clusters. The rationale for this split analysis was that due to the retinotopic processing left-hemispheric alpha generators may have entrained with the 12-Hz stimulus while right-hemispheric generators may have entrained with the 10-Hz stimulus.

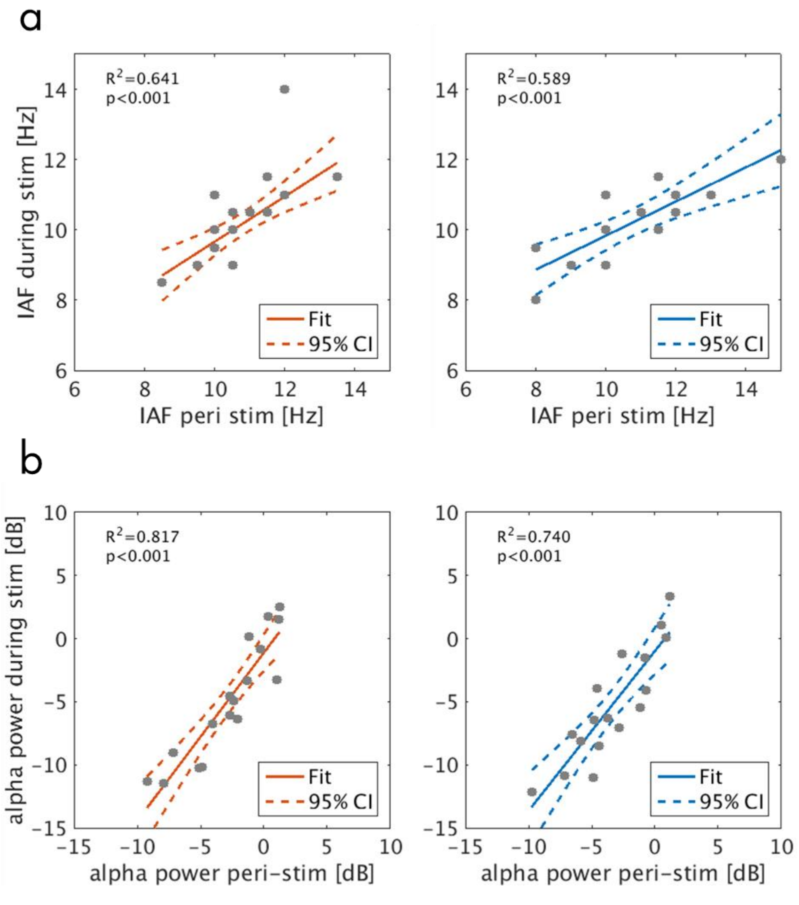


**Figure S5** Linear regression: Individual alpha frequency (IAF) during constant-frequency stimulation (y-axis) as a function of IAF peri-stimulation (x-axis). Left (red) plot = data from right electrode cluster, i.e. contralateral to the 10-Hz stimulus. Right (blue) plot = data from left electrode cluster, i.e. contralateral to the 12-Hz stimulus. Grey dots represent participants. Coloured lines depict a straight line fit and its confidence interval (dashed lines). Goodness-of-fit of the linear model provided as $R^2$ along with corresponding *P*-Value. (**b**) Same as in (a) but for spectral power (in dB) at IAF. Compare with *Figure 5*.

As illustrated in *Figure S5* and detailed in *Table S1,* IAF and IAP showed systematic linear dependencies between measures taken peri- and during stimulation. Crucially, direct comparisons of regression slopes ($\beta_1$) derived from left and right hemispheres did not reveal differences for IAF ($\Delta\beta_1$ = 0.191, *CI* = [-0.421 0.516]).

Due to the slightly different approaches we abstained from a direct statistical comparison of regression results with frequency-varying conditions but note that IAF regression slopes reported here and their counterparts were of similar magnitude (compare IAF $\beta_1$ in *Table 1* and *Table S1*). This suggests that strictly rhythmic stimulation did not have a distinct influence on the intrinsic alpha rhythm either.

**Table S1** Parameters of robust linear regressions for the constant frequency stimulation condition, separated by hemisphere/stimulation frequency. Predictor = alpha measure (IAF, IAP) peri-stimulation; outcome = alpha measure during stimulation. 95% confidence intervals (*CI*) were bootstrapped (*N* = 10000) using the bias-corrected-and-accelerated method.

| | *Hemisphere* | $R^2$ | $\beta_0$ | *95% CI* | $\beta_1$ | *95% CI* |
|---|---|---|---|---|---|---|
| IAF | Left (12 Hz) | 0.589 | -0.013 | [-0.328 0.311] | 0.794 | [0.504 1.220]* |
| | Right (10 Hz) | 0.641 | -0.180 | [-0.347 0.255] | 0.603 | [0.262 0.818]* |
| IAP | Left (12 Hz) | 0.740 | -0.988 | [-3.074 0.557] | 1.263 | [0.922 1.641]* |
| | Right (10 Hz) | 0.817 | -1.130 | [-3.697 0.088] | 1.328 | [0.854 1.984]* |

IAF – individual alpha frequency; IAP – individual alpha power; $R^2$ – goodness of fit; $\beta_0$ – intercept; $\beta_1$ – slope; * denotes significance
